# Mapping consistent, reproducible, and transcriptionally relevant functional connectome hubs of the human brain

**DOI:** 10.1101/2021.11.29.470494

**Authors:** Zhilei Xu, Mingrui Xia, Xindi Wang, Xuhong Liao, Tengda Zhao, Yong He

## Abstract

Human brain connectomes include sets of densely connected regions, known as connectome hubs, which play a vital role in understanding global brain communication, cognitive processing, and brain disorders. However, the consistency and reproducibility of functional connectome hubs’ anatomical localization have not been established to date and the genetic signatures underlying robust connectome hubs remain unknown. Here, we conduct the first worldwide, harmonized meta-connectomic analysis by pooling resting-state functional MRI data of 5,212 healthy young adults across 61 independent cohorts. We identify highly consistent and reproducible functional connectome hubs in heteromodal and unimodal regions both across cohorts and across individuals. These connectome hubs show heterogeneous connectivity profiles and are critical for both intra- and inter-network communications. Using post-mortem gene expression data, we show that these connectome hubs have a spatiotemporally distinctive transcriptomic pattern dominated by genes involved in the neuropeptide signaling pathway, neurodevelopmental processes, and metabolic processes. These results highlight the robustness of macroscopic connectome hubs and their potential cellular and molecular underpinnings.

## Introduction

Functional connectome mapping studies have identified sets of densely connected regions in large-scale human brain networks, which are known as hubs^1^. Connectome hubs play a crucial role in global brain communication^1, 2^ and support a broad range of cognitive processing, such as working memory^3^ and semantic processing^4^. Growing evidence suggests that these highly connected brain hubs are preferentially targeted by many neuropsychiatric disorders^5–8^, which provides critical clues for understanding the biological mechanisms of disorders and establishing biomarkers for disease diagnosis^8, 9^ and treatment evaluation^10, 1, 2, 11, 12^ (for reviews).

Despite such importance, there is considerable inconsistency in anatomical locations of functional connectome hubs among existing studies. For example, components of the defaultmode network (DMN) have been frequently reported as connectome hubs, yet the spatial pattern is highly variable across studies. In particular, several studies have shown highly connected hubs in the lateral parietal regions of the DMN^7, 8, 13, 14^, whereas others have reported midline structures of the DMN^15–19^. Several works have identified primary sensorimotor and visual regions as connectome hubs^13, 14, 16–19^, yet others did not replicate these findings^7, 8, 15^. Subcortical regions, such as the thalamus and amygdala, have also been inconsistently reported as hubs^8, 15, 16, 18^ and non-hubs^7, 13, 14, 17, 19^ Thus, the consistency and reproducibility of functional connectome hubs have been difficult to establish to date, which can be attributed to inadequate sample size and differences in imaging scanner, imaging protocol, data processing, and connectome analysis strategies. Here, we aimed to establish a harmonized meta-analysis model to identify robust functional connectome hubs in healthy young adults by combining multiple cohorts with uniform protocols for data quality assurance, image processing, and connectome analyses.

Once the robust connectome hubs are identified, we will further examine their genetic signatures. It has been well demonstrated that the connectome architecture of the human brain is inheritable, such as functional connectivity of the DMN^20^ and the cost-efficiency optimization^21^. Moreover, the functional connectomes can be regulated by genotypic variation both during rest^22^ and in cognitive tasks^23^, especially involving the DMN^22, 23^ and frontoparietal network (FPN)^23^. Growing evidence also suggests spatial correspondence between transcriptomic profiles and connectome architectures^24–26, 27^ (for review). Thus, we reasoned that the robust macroscopic connectome hubs could be associated with microscopic genetic signatures. Elucidating these genetic signatures will substantially benefit our understanding of how connectome hubs emerge in development, function in complex cognition, and are involved in disease.

To address these issues, we provide the first worldwide harmonized meta-connectomic analysis of functional brain hubs by pooling a large-sample resting-state functional MRI (rsfMRI) dataset of 5,212 healthy young adults (aged 18-36 years) across 61 cohorts. Fig 1 illustrates the sample size and age ranges of each cohort. To uncover the genetic signatures underlying these connectome hubs, we conducted machine learning approaches to distinguish connectome hubs from non-hubs using transcriptomic data from the Allen Human Brain Atlas (AHBA) and explored their developmental evolutions using the BrainSpan Atlas.

**Fig 1.**
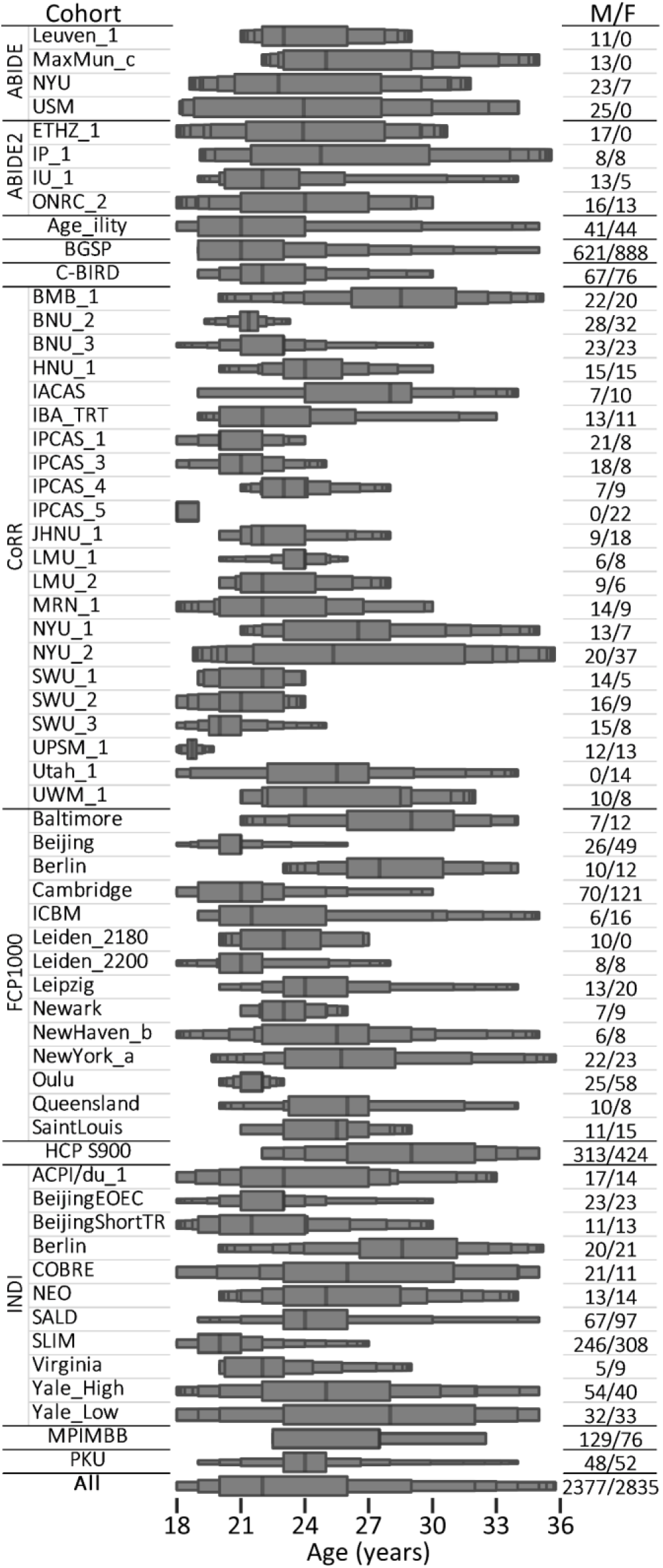
Enhanced box plot of the age ranges of each cohort. M/F, males/femals.

## Results

### Identifying consistent connectome hubs using a harmonized meta-analysis model

Prior to the meta-analysis, we constructed a voxelwise functional connectome matrix for each individual by computing the Pearson’s correlation coefficient between preprocessed rsfMRI time series of all pairs of gray matter voxels (47,619 voxels). Then, the functional connectivity strength (FCS) of each voxel was computed as the sum of connection weights between the given voxel and all the other voxels. This resultant FCS map was further normalized with respect to its mean and standard deviation across voxels^7^. For each cohort, we performed a general linear model on these normalized FCS maps to reduce age and gender effects. As a result, we obtained a mean FCS map and its corresponding variance map for each cohort that were used for subsequent meta-analyses.

To identify the most consistent connectome hubs, we conducted a voxelwise random-effects meta-analysis on the mean and variance FCS maps of the 61 cohorts. Such an analysis addressed the across-cohort heterogeneity of functional connectomes, resulting in a robust FCS pattern and its corresponding standard error (SE) map (Fig 2A). Then, we identified consistent connectome hubs whose FCS values were significantly *(p* < 0.001, cluster size > 200 mm^3^) higher than the global mean (i.e., zero) using a voxelwise *Z* value map computed by dividing the FCS map by the SE map. To determine the significance levels of these observed *Z* values, a nonparametric permutation test^28^ with 10,000 iterations was performed. Finally, we estimated voxelwise effect sizes using Cohen’s *d* metric computed by dividing the *Z* value map by the square root of the cohort number (Fig 2B, left). According to prior brain network parcellations^29, 30^, these identified hub voxels (15,461 voxels) were spatially distributed in multiple brain networks, including the DMN (27.5%), dorsal attention network (DAN) (16.5%), FPN (15.9%), ventral attention network (VAN) (15.6%), somatomotor network (SMN) (14.4%), and visual network (VIS) (9.9%) (Fig 2B, right). Using a local maxima localization procedure, we identified 35 robust brain hubs across 61 cohorts (Fig 2B, left; Table 1), involving various heteromodal and unimodal areas. Specifically, the most robust findings resided in several lateral parietal regions, including the bilateral ventral postcentral gyrus, supramarginal gyrus, and angular gyrus.

**Fig 2.**
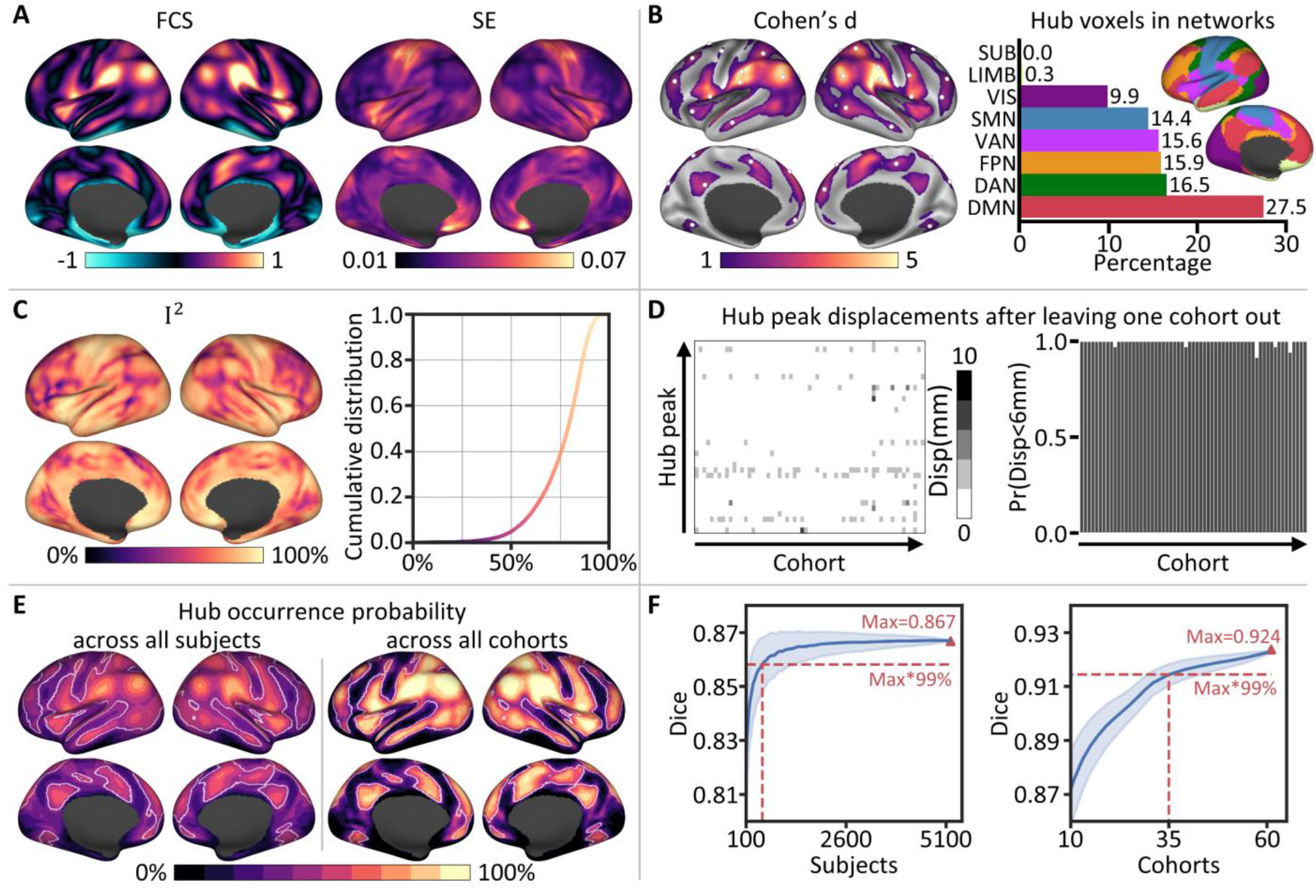
Highly consistent and reproducible functional connectome hubs. **A** Robust FCS pattern and its corresponding variance (standard error, SE) map estimated using a harmonized voxelwise random-effects meta-analysis across 61 cohorts. **B** Left: The most consistent functional connectome hubs *(p* < 0.001, cluster size > 200 mm^3^); white spheres represent hub peaks. Right: Hub voxel distribution in eight large-scale brain networks; insets, the seven large- scale cortical networks^29^ were rendered on the left hemisphere. SUB, subcortical network; LIMB, limbic network. **C** Left: Heterogeneity measurement *I*^2^ estimated through the randomeffects meta-analysis. Right: Cumulative distribution function plot of *I^2^.* **D** Left: Heatmap of displacements of the 35 hub peaks after leaving one cohort out. Right: Bar plot of the probability across the 35 hub peaks whose displacement was less than 6 mm after leaving one cohort out. **E** Hub occurrence probability map across all subjects (left) and all cohorts (right). White lines delineate boundaries of the identified hubs in **B. F** Dice’s coefficient of the identified hubs in **B** compared with the top *N* (voxel number of the identified hubs in **B**) voxels with the highest hub occurrence probability values across randomly selected subjects (left) and randomly selected cohorts (right). Blue shading represents the standard deviation across 2,000 random selections.

**Table 1.** Highly consistent functional connectome hubs.

| No. | Hub | Location | MNI coordinates |  |  | Cohen's *d* | FCS | SE |
| --- | --- | --- | --- | --- | --- | --- | --- | --- |
|  |  |  | x | y | z |  |  |  |
| 1 | Right PFt | PFt (superoanterior BA* 40) | 60 | -21 | 45 | 6.267 | 1.072 | 0.022 |
| 2 | Left PFt | PFt (superoanterior BA 40) | -60 | -24 | 36 | 6.151 | 0.949 | 0.020 |
| 3 | Right PF | PF (posterior BA 40) | 60 | -27 | 24 | 5.785 | 1.239 | 0.027 |
| 4 | Left SCEF | Supplementary and cingulate eye field | 0 | 0 | 51 | 5.635 | 1.000 | 0.023 |
| 5 | Left PGi | PGi (inferior BA 39) | -51 | -66 | 30 | 5.168 | 1.075 | 0.027 |
| 6 | Left PFop | PF opercular (inferoanterior BA 40) | -63 | -27 | 18 | 5.160 | 1.095 | 0.027 |
| 7 | Left 43 | Area 43 | -57 | 3 | 3 | 4.927 | 1.114 | 0.029 |
| 8 | Right 6r | Rostral area 6 | 57 | 6 | 0 | 4.916 | 1.184 | 0.031 |
| 9 | Right PGi | PGi (inferior BA 39) | 54 | -60 | 30 | 4.739 | 1.007 | 0.027 |
| 10 | Right 8BL | Area 8B lateral | 21 | 36 | 51 | 4.655 | 0.713 | 0.020 |
| 11 | Right 7PC | Area 7PC | 36 | -45 | 54 | 4.414 | 0.712 | 0.021 |
| 12 | Left 9p | Area 9 posterior | -15 | 45 | 45 | 4.199 | 0.639 | 0.019 |
| 13 | Right 6v | Ventral area 6 | 54 | 9 | 33 | 4.037 | 0.766 | 0.024 |
| 14 | Left 8Av | Ventral area 8A | -39 | 18 | 48 | 3.990 | 0.561 | 0.018 |
| 15 | Left AIP | Anterior intra-parietal area | -33 | -45 | 45 | 3.474 | 0.567 | 0.021 |
| 16 | Right FST | Fundus of the superior temporal area | 54 | -60 | 0 | 3.156 | 0.729 | 0.030 |
| 17 | Right 9m | Area 9 middle | 3 | 54 | 24 | 3.128 | 0.609 | 0.025 |
| 18 | Left 31pv | Area 31p ventral | -3 | -51 | 33 | 3.049 | 0.784 | 0.033 |
| 19 | Right VIP | Ventral intra-parietal complex | 18 | -63 | 57 | 2.984 | 0.572 | 0.025 |
| 20 | Right 6a | Area 6 anterior | 33 | 3 | 63 | 2.975 | 0.454 | 0.020 |
| 21 | Left FOP4 | Frontal opercular area 4 | -33 | 21 | 6 | 2.858 | 0.828 | 0.037 |
| 22 | Right 5mv | Area 5m ventral | 12 | -30 | 45 | 2.822 | 0.701 | 0.032 |
| 23 | Right 46 | Area 46 | 36 | 42 | 30 | 2.779 | 0.656 | 0.030 |
| 24 | Left 10v | Area 10v | 0 | 57 | -9 | 2.769 | 0.731 | 0.034 |
| 25 | Left p9-46v | Area posterior 9-46v | -42 | 36 | 27 | 2.591 | 0.561 | 0.028 |
| 26 | Left V3A | Area V3A | -15 | -90 | 33 | 2.575 | 0.684 | 0.034 |
| 27 | Left TE1a | Area TE1 anterior | -63 | -15 | -15 | 2.527 | 0.595 | 0.030 |
| 28 | Right TE1a | Area TE1 anterior | 60 | -9 | -21 | 2.494 | 0.580 | 0.030 |
| 29 | Right IFSa | Anterior inferior frontal sulcus | 48 | 39 | 12 | 2.468 | 0.480 | 0.025 |
| 30 | Left 7Am | Medial area 7A | -12 | -60 | 60 | 2.461 | 0.475 | 0.025 |
| 31 | Right V3A | Area V3A | 18 | -87 | 36 | 2.442 | 0.645 | 0.034 |
| 32 | Right V4 | Fourth visual area | 24 | -63 | -9 | 2.339 | 0.446 | 0.024 |
| 33 | Left 6a | Area 6 anterior | -24 | 3 | 63 | 2.317 | 0.331 | 0.018 |
| 34 | Left VMV1 | Ventromedial visual area 1 | -18 | -60 | -6 | 1.937 | 0.397 | 0.026 |
| 35 | Left FEF | Frontal eye fields | -45 | -9 | 57 | 1.412 | 0.640 | 0.058 |

\* BA, Brodmann area.

### The identified connectome hubs are reproducible across cohorts and individuals

During identifying the above highly consistent connectome hubs, the random-effects metaanalysis revealed high heterogeneity of FCS across cohorts (Fig 2C, left). The cumulative distribution function plot shows more than 95% voxels with *I^2^* (heterogeneity score) exceeding 50% (Fig 2C, right), indicating high heterogeneity across cohorts in almost all brain areas (see also Fig S1). To determine whether the connectome hubs identified here are dominated by certain cohorts or are reproduced across-subject/cohort, we performed a leave-one-cohort-out validation analysis and an across-subject/cohort conjunction analysis.

#### Leave-one-cohort-out validation analysis

We repeated the above harmonized meta-analysis hub identification procedure after leaving one cohort out at a time. Comparing the identified hubs using all cohorts (Fig 2B) with those after leaving one cohort out obtained extremely high Dice’s coefficients (*mean±sd:* 0.990±0.006; range: 0.966-0.997). For hub peaks, leaving one cohort out resulted in very few displacements (mostly fewer than 6 mm, Fig 2D). Thus, connectome hubs identified using the 61 cohorts were not dominated by specific cohorts.

#### Across-subject/cohort conjunction analysis

We defined the top *N* (*N* = 15,461, which is the voxel number of hubs in Fig 2B) voxels with the highest FCS values of a subject or a cohort as connectome hubs for that subject or that cohort. Then, for each voxel, we assessed hub occurrence probability values across subjects and cohorts. The identified hubs using all cohorts were highly overlapped with the top *N* voxels with the highest hub occurrence probability values both across all subjects and across all cohorts, indicated by a high Dice’s coefficient *(Dice* = 0.867, Fig 2E, left; *Dice* = 0.924, Fig 2E, right). When the identified hubs using all cohorts were compared with the top *N* voxels with the highest hub occurrence probability values across randomly selected subjects or across randomly selected cohorts, the Dice’s coefficient approached 99% of its maximum value after exceeding 510 subjects (Fig 2F, left) and 35 cohorts (Fig 2F, right), respectively. This indicated that the identified connectome hubs were highly reproducible both across cohorts and across individuals.

Validation analysis demonstrated that the above results did not depend on analysis parameters, such as the connection threshold (Fig S2 and S3), and were not driven by the size of the brain network to which they belong^31^ (Fig S4), suggesting the robustness of our main findings.

### Connectome hubs have heterogeneous functional connectivity profiles

Next, we further examined whether these robust brain hubs (Fig 2B and Table 1) have distinctive functional connectivity profiles that represent their unique roles in network communication. To gain detailed and robust functional connectivity profiles of each hub region, we conducted a seed-to-whole-brain connectivity meta-analysis in a harmonized protocol again. For each of the 35 hub regions, we obtained an estimated Cohen’s *d* effect size map that characterizes the robust whole-brain connectivity pattern relevant to the seed region across the 61 cohorts (Fig 3A). We then divided the connectivity map of each hub into eight brain networks according to prior parcellations^29, 30^, resulting in an 8×35 connectivity matrix with each column representing the voxel percentage of each of the eight networks connected with a hub.

**Fig 3.**
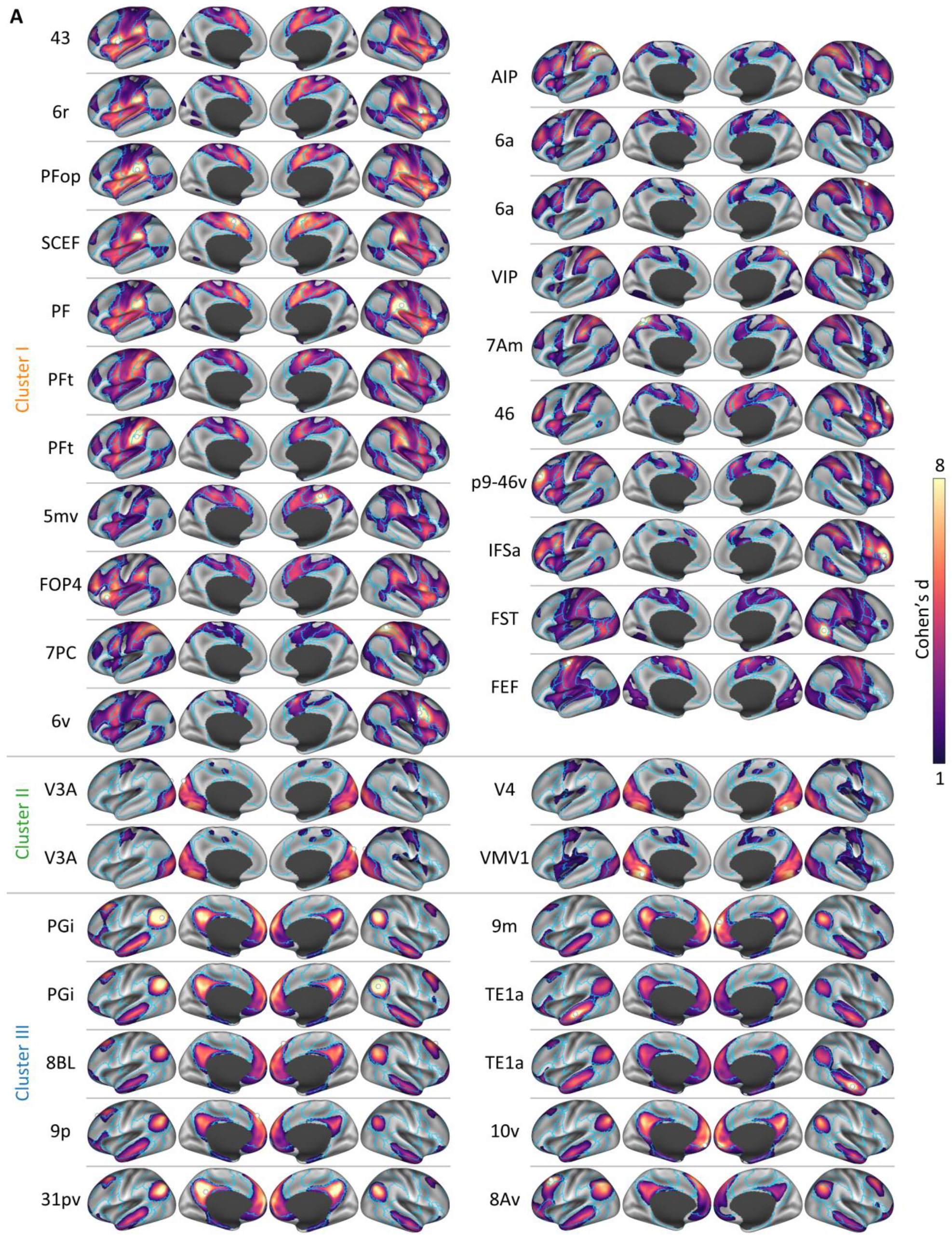
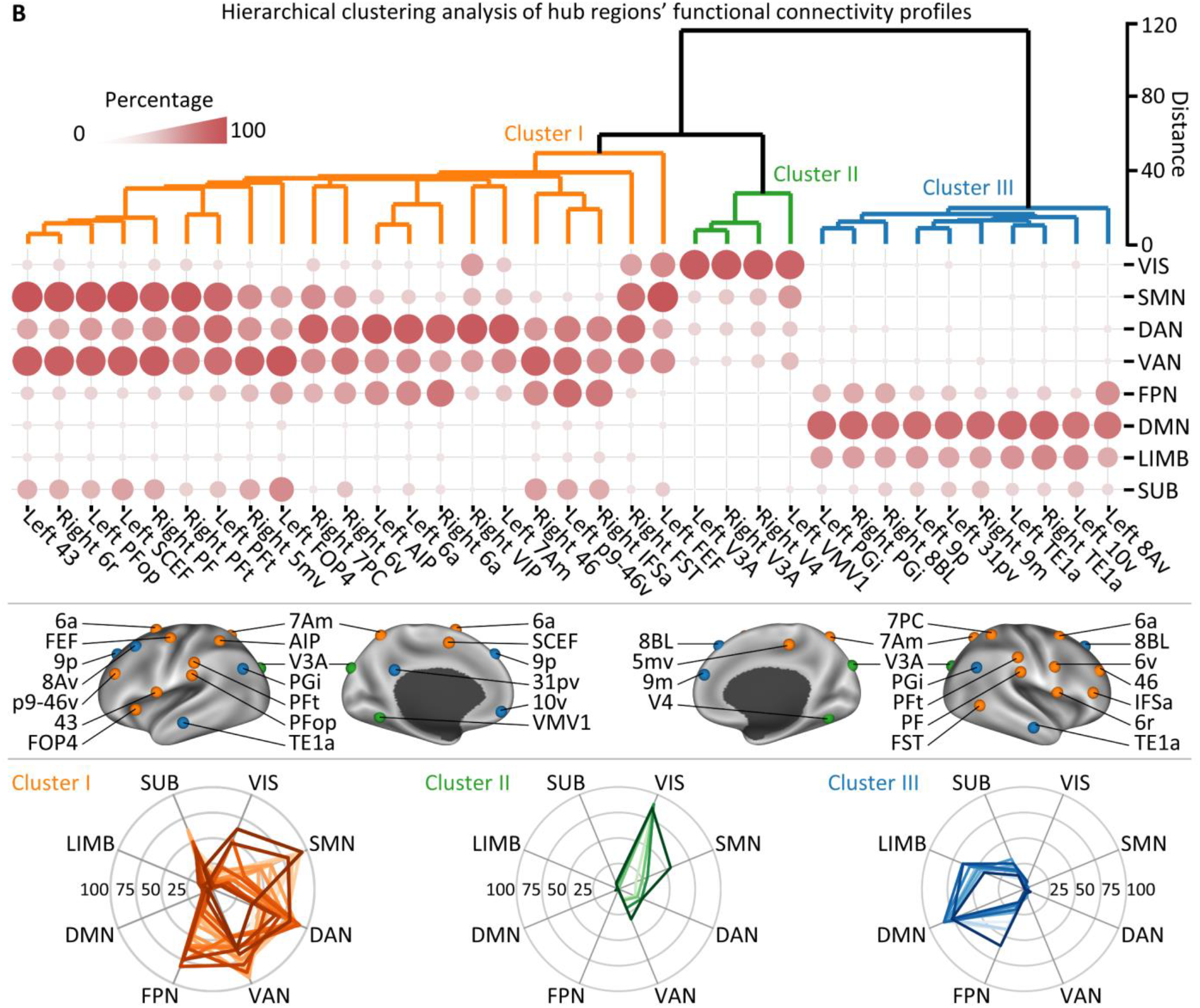
Functional connectivity profiles of connectome hubs. **A** Functional connectivity profiles of the 35 hubs. White spheres represent hub seeds. Blue lines delineate boundaries of the seven cortical networks shown in Fig 2B. **B** Top: Dendrogram derived by hierarchical clustering of the connectivity percentage matrix. Middle: The 35 hubs were rendered using three different colors according to the hierarchical clustering solution. Bottom: Radar charts showing heterogeneous connectivity profiles of the three hub clusters.

Hierarchical clustering analysis on the connectivity matrix clearly divided the 35 hubs into three clusters (Fig 3B). Cluster I consists of 21 hubs that are primarily connected with extensive areas in the DAN, VAN, FPN, and SMN (orange, Fig 3B). Cluster II consists of four hubs that are densely connected with VIS (green, Fig 3B). Cluster III consists of 10 hubs that have robust connections with the DMN and LIMB (blue, Fig 3B). Of particular interest is that within Cluster III, a left posterior middle frontal hub called ventral area 8A (8Av) shows a distinctive connectivity profile in contrast to the other nine hubs, manifested as having robust connections with bilateral frontal FPN regions (Fig 3A and Fig S5). This imply that the left 8Av hub is a key connector between the DMN and FPN, which can be supported by the recent finding of a control-default connector located in the posterior middle frontal gyrus^32^. Although both Cluster I and III hubs are connected with subcortical structure, they are connected with different subcortical nuclei (Fig 3B and Fig S6). Finally, whereas all hubs possess dense intranetwork connections, most also retain significant internetwork connections (Fig S7), which preserves efficient communication across the whole brain network feasible.

### Transcriptomic data distinguishes connectome hubs from non-hubs

A supervised machine learning classifier based on XGBoost^33^ and 10,027 genes’ transcriptomic data from the AHBA^34^ was trained to distinguish connectome hubs from non-hubs (Fig 4A). The sensitivity, specificity, and accuracy rate of the XGBoost classifier were stably estimated by repeating the training and testing procedure 1,000 times. This classifier performed better than chance in all 1,000 repetitions and achieved an overall accuracy rate of 65.3% (Fig 4B). In crossvalidation, connectome hubs and non-hubs were classified with a sensitivity of 71.1% and specificity of 63.4%, respectively. The testing procedure yielded a comparable sensitivity of 69.7% and specificity of 62.0%. After training the classifier, each gene’s contribution to the optimal prediction model was determined. We noted that some key genes contributed two or three orders of magnitude more than other genes (Fig 4C). The contributions of the top 300 mostly contributed key genes were consistent between the first 500 repetitions and the second 500 repetitions (Pearson’s *r* = 0.958,*p* < 10^-6^, Fig 4D), suggesting a high reproducibility.

**Fig 4.**
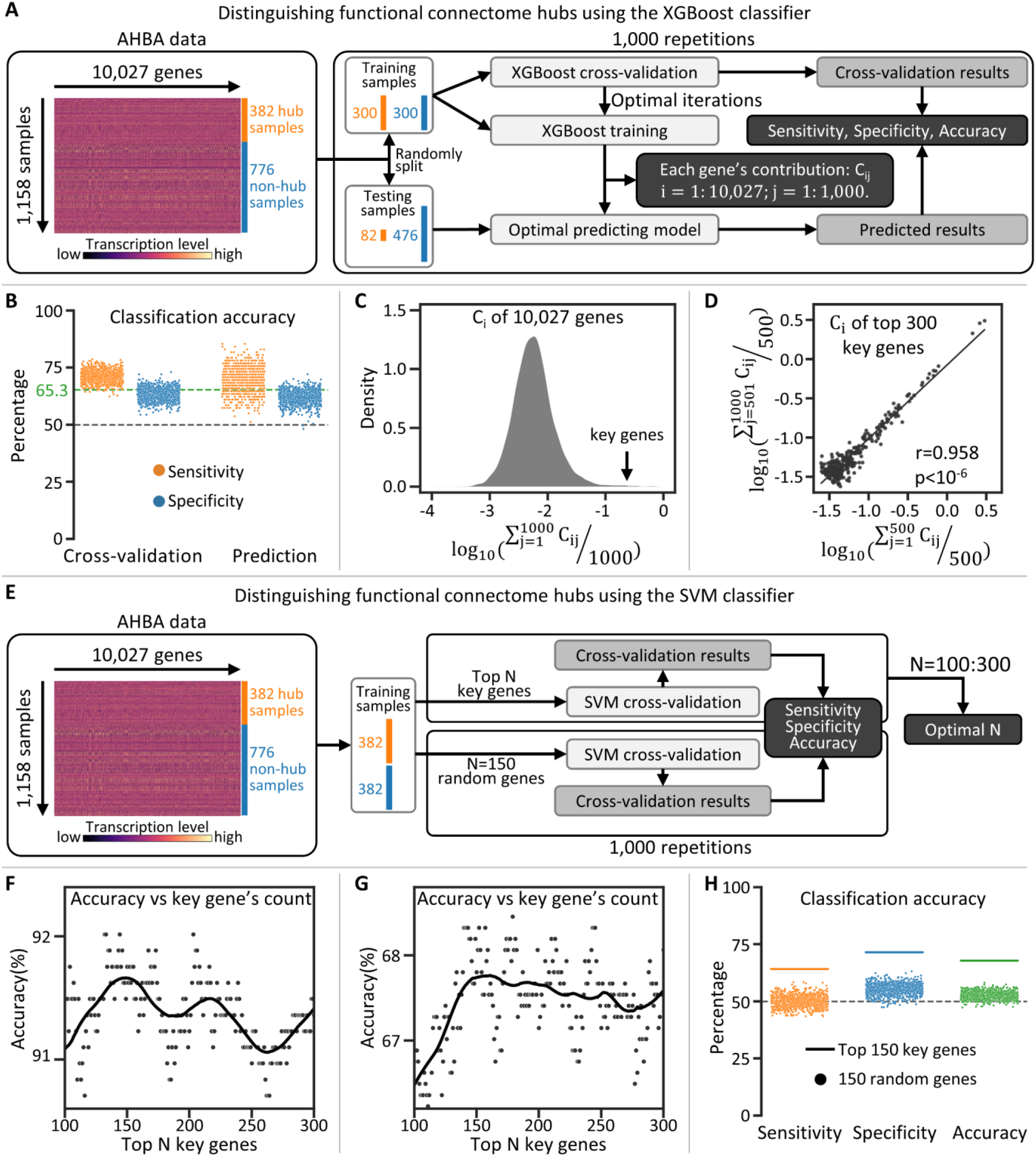
Transcriptomic data distinguishes connectome hubs from non-hubs. **A** Schematic diagram of using the XGBoost model to classify brain samples as a hub or non-hub. **B** Performance of the XGBoost classifier. Each dot represents one repetition in **A**. The horizontal gray dashed line represents the chance level accuracy rate (50%). The horizontal green dashed line represents the average accuracy rate of the XGBoost classifier across 1,000 repetitions. **C** Density plot of 10,027 genes’ logarithmic average contributions across 1,000 repetitions to the XGBoost classifier. Genes with the greatest contributions were regarded as key genes. **D** Regression plot of the logarithmic average contributions of the top 300 key genes across the first 500 repetitions versus those across the second 500 repetitions. Each dot represents one gene. **E** Schematic diagram of using the SVM model to classify brain samples as a hub or non-hub. **F and G** Accuracy rate of the SVM classifier versus the count of key genes used to distinguish 382 hub samples from 382 non-hub samples with the highest rate (**F**) or lowest rate (**G**) to be correctly classified by the XGBoost classifier. Each dot represents one SVM classifier. Black curves were estimated by locally weighted regression. **H** Performance of the SVM classifier. Horizontal lines correspond to the SVM classifier trained using top 150 key genes in **G**. Each dot represents one repetition using 150 randomly selected genes in **E**. The horizontal gray dashed line represents the chance level accuracy rate (50%).

To exclude the XGBoost model’s potential bias relating to the mostly contributed key genes, we replicated the above classification results using another machine learning model based on the support vector machine (SVM) that was trained using only the top *N* key genes with the greatest contributions to the XGBoost classifier (Fig 4E). Because no data were available to determine how many key genes were sufficient to train an SVM classifier, we examined the count *N* from 100 to 300. The SVM classifier achieved a very high peak accuracy rate of 91.8% with approximately the top 150 key genes in the easiest classification task (Fig 4F) and also achieved a reasonable peak accuracy rate of 67.8% with approximately the top 150 key genes even in the most difficult classification task (Fig 4G). By contrast, SVM classifiers trained using 150 randomly selected genes performed worse than that using the top 150 key genes in all 1,000 repetitions (Fig 4H). Thus, these robust connectome hubs were significantly associated with a transcriptomic pattern dominated by approximately 150 key genes (Table S1).

### Connectome hubs have a spatiotemporally distinctive transcriptomic pattern

Gene Ontology (GO) enrichment analysis using GOrilla^35^ demonstrated that the above 150 key genes were mostly enriched in the neuropeptide signaling pathway *(fold enrichment (FE)* = 8.9, uncorrected *p* = 1.2×10^-5^, Table S2). GO enrichment analysis using the ranked 10,027 genes according to their contributions to the XGBoost classifier also confirmed the most enriched GO term of the neuropeptide signaling pathway (*FE* = 5.7, uncorrected *p* < 10^-6^, Table S3). The ranked 10,027 genes were also associated with the developmental process (*FE* = 1.2), cellular developmental process (*FE* = 1.3), anatomical structure development (*FE* = 1.3), and neuron projection arborization (*FE* = 13.7) (uncorrected*ps* < 5.5×10^-4^, Table S3). We speculated that connectome hubs have a distinctive transcriptomic pattern of neurodevelopmental processes in contrast to non-hubs.

We repeated the GO enrichment analysis of the above 150 key genes using DAVID^36, 37^ In addition to the mostly enriched GO term of the neuropeptide signaling pathway (*FE* = 8.7, uncorrected *p* = 5.8×10^-4^), there were 10 GO terms associated with metabolic process, such as the positive regulation of cellular metabolic process (*FE* = 1.4, uncorrected *p* = 0.031, Table S4). Disease association analysis demonstrated metabolic disease associated with the greatest number of key genes (60 genes, *FE* = 1.2, uncorrected *p* = 0.094, Table S5). Accordingly, it is rational to speculate that connectome hubs have a distinctive transcriptomic pattern of metabolic processes in contrast to non-hubs.

To confirm the above two speculations of GO enrichment analysis results, we examined transcription level differences between hub and non-hub regions for genes previously implicated in key neurodevelopmental processes^38^ (Table S6) and main neuronal metabolic pathways^39^ (oxidative phosphorylation^40^ and aerobic glycolysis^41^, Table S7). Permutation tests revealed hub regions with significantly higher transcription levels for genes associated with dendrite development, synapse development, and aerobic glycolysis than non-hub regions (one-sided Wilcoxon rank-sum tests, Bonferroni-corrected *ps* ≤ 0.032, Fig 5A). In addition, hub regions had a weak trend of lower transcription levels for genes associated with axon development, myelination, neuron migration, and oxidative phosphorylation (Fig 5A). These transcription level differences were consistent with our speculations of GO enrichment analysis results.

**Fig 5.**
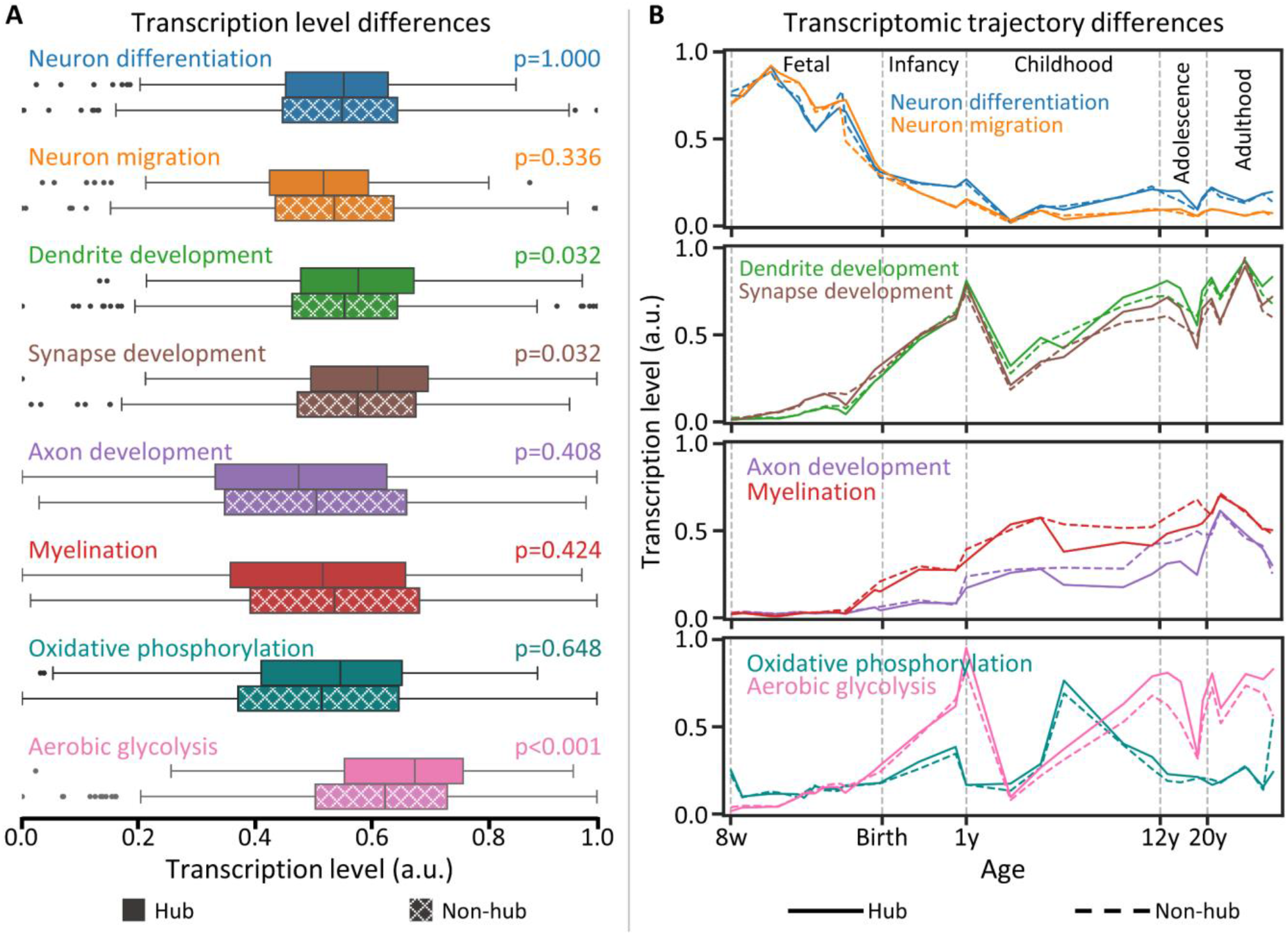
Connectome hubs have a spatiotemporally distinctive transcriptomic pattern. **A** Transcription level differences between hub samples (n=382) and non-hub samples (n=776) for genes associated with key neurodevelopmental processes^38^ and main neuronal metabolic pathways^39^. Boxplot edges, gray lines, and whiskers and dots depict the 25th and 75th percentiles, median, and extreme nonoutlier and outlier values, respectively. Significance of onesided Wilcoxon rank-sum tests were determined by 1,000 permutation tests and were labeled with Bonferroni-corrected *p* values. **B** Transcriptomic trajectory differences between hub and non-hub regions for genes involved in key neurodevelopmental processe^38^ and main neuronal metabolic pathways^39^. w, post-conceptional week; y, postnatal year; a.u., arbitrary unit.

These above transcriptomic results were derived from the AHBA, an adult transcriptomic dataset. To explore their developmental evolutions, we inspected transcriptomic trajectory differences between hub and non-hub regions using the BrainSpan Atlas^42^. We observed pronounced diverging transcriptomic trajectories between hub and non-hub regions in these key neurodevelopmental processes and main neuronal metabolic pathways (Fig 5B and Fig S8). For neuron migration, the transcription level in hub regions is higher than that in non-hub regions from the mid-fetal period to early infancy. For dendrite, synapse, axon development, and myelination, transcriptomic trajectories of hub regions apparently diverge from those of nonhubs during childhood and adolescence, during which hub regions have higher transcription levels for dendrite and synapse development but lower transcription levels for axon development and myelination. These results are in agreement with the observation of primary somatosensory, auditory, and visual (V1/V2) cortices with lower synapse density but higher myelination than the prefrontal area^43, 44^. Mmoreover, hub regions have higher transcription levels than non-hub regions for aerobic glycolysis since the early childhood period and for oxidative phosphorylation during childhood and adolescence. These transcriptome analyses achieved convergent results between the AHBA and BrainSpan Atlas.

Together, functional connectome hubs have a spatiotemporally distinctive transcriptomic pattern in contrast to non-hubs, which is dominated by genes involved in the neuropeptide signaling pathway, neurodevelopmental processes, and metabolic processes.

### Connectome hubs have more intricate fiber configuration and higher metabolic rate

Growing evidence has suggested a striking spatial correspondence between transcriptomic profile and structural connectivity in the human brain^27^. We speculated that the above microscale transcriptomic differences between hub and non-hub regions in key neurodevelopmental processes may result in macroscale structural connectivity profile differences. Using a fiber length profiling dataset^45^, we observed that hub regions possess more fibers with a length exceeding 40 mm but less fibers with a length shoter than 40 mm (one-sided Wilcoxon rank-sum tests, Bonferroni-corrected *ps* ≤ 0.008, Fig 6). That is, hub regions have more short, medium, and long fibers, whereas non-hub regions have more very short (< 40 mm) fibers, suggesting a more intricate fiber configuration in hub regions.

**Fig 6.**
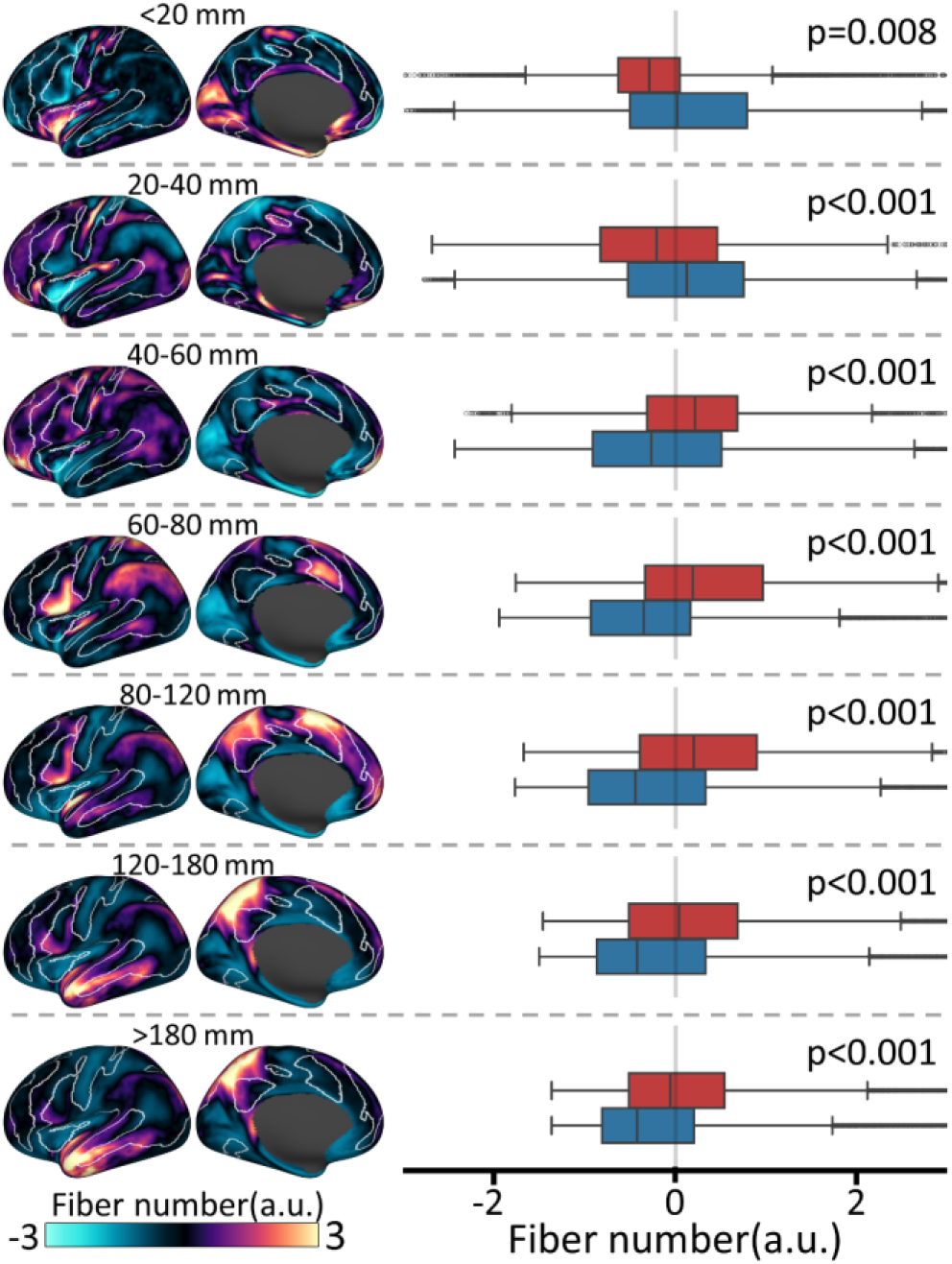
Connectome hubs have more intricate fiber configuration. Left: Fiber number for different fiber length bins was derived from a fiber length profiling dataset^45^. To save space, we only presented data of the left hemisphere and used data of both hemispheres in boxplots and statistics. White lines delineate boundaries of the identified hubs in Fig 2B. Right: Fiber number difference between connectome hubs (red, n=25,944) and non-hubs (blue, n=33,195). Boxplot edges, gray lines, and whiskers and dots depict the 25th and 75th percentiles, median, and extreme nonoutlier and outlier fiber number values, respectively. Significance of one-sided Wilcoxon rank-sum tests were determined by 1,000 permutation tests and were labeled with Bonferroni-corrected *p* values. a.u., arbitrary unit.

The above transcriptome analyses have shown a higher transcription level of oxidative phosphorylation and aerobic glycolysis in hub regions than in non-hubs. We validated these observations using a metabolism dataset derived from positron emission tomography^46^ and found that hub regions not only have a higher metabolic rate than non-hubs in oxidative phosphorylation (indicated by the cerebral metabolic rate for oxygen) and aerobic glycolysis (indicated by the glycolytic index), but also have more blood supply (indicated by the cerebral blood flow) (one-sided Wilcoxon rank-sum tests, Bonferroni-corrected *ps* < 0.001, Fig 7). This is in agreement with prior observations of a tight coupling between FCS and blood supply^1, 47^

**Fig 7.**
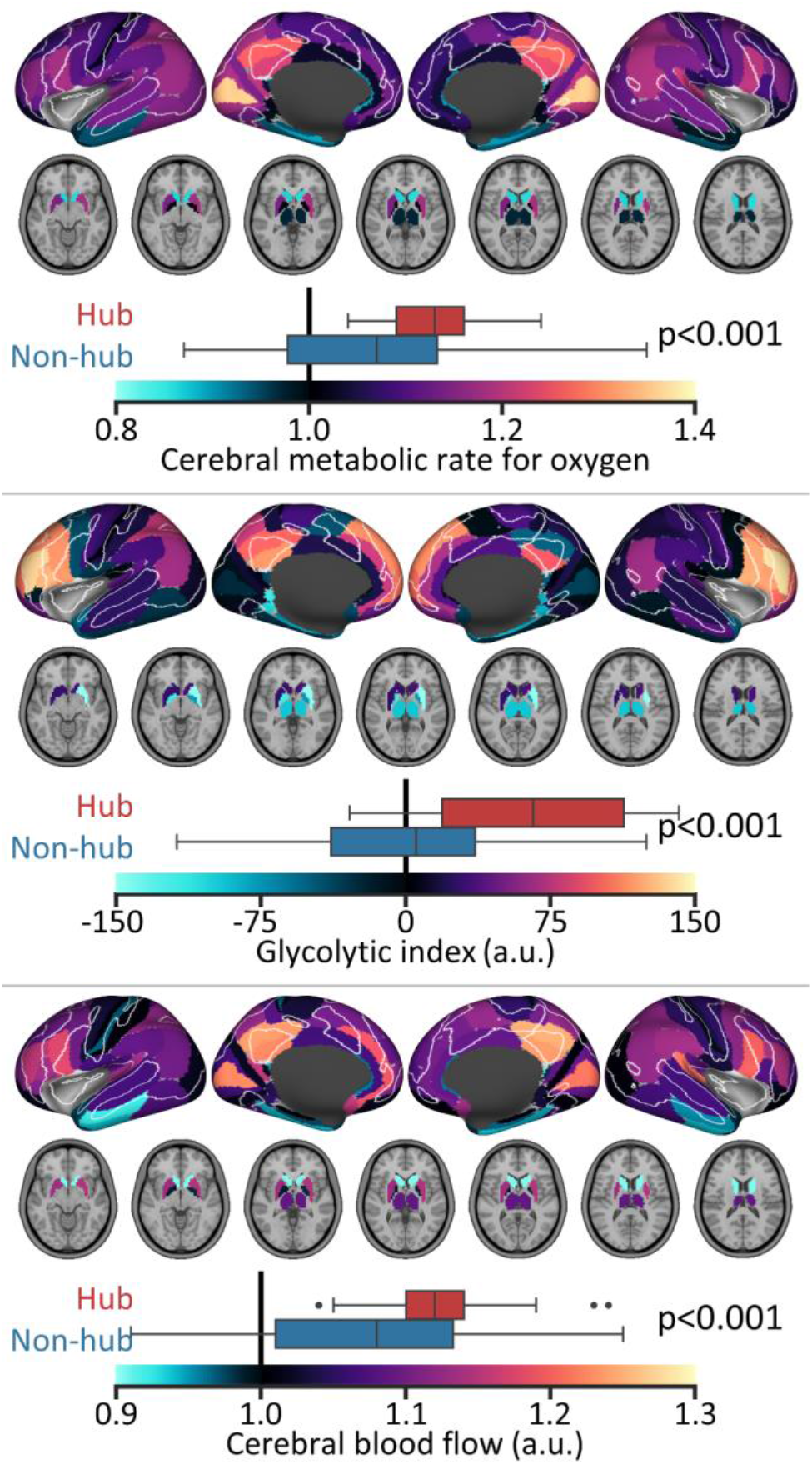
Connectome hubs have higher metabolic rate. The cerebral metabolic rate for oxygen, glycolytic index, and cerebral blood flow of 82 Brodmann areas and seven subcortical structures were provided by a prior study^46^. White lines delineate boundaries of the identified hubs in Fig 2B. Boxplot edges, gray lines, and whiskers and dots depict the 25th and 75th percentiles, median, and extreme nonoutlier and outlier metabolic measurement values, respectively. Brodmann areas with more than 50% vertices or subcortical structures with more than 50% voxels identified as hubs were regarded as hub regions (n=29), vice versa as non-hub regions (n=60). Significance of one-sided Wilcoxon rank-sum tests was determined by 1,000 permutation tests and were labeled with Bonferroni-corrected*p* values. a.u., arbitrary unit.

## Discussion

Using a worldwide harmonized meta-connectomic analysis of 5,212 healthy young adults across 61 cohorts, we provided the first description of highly consistent and reproducible functional connectome hubs in the resting human brain. Using transcriptomic data from the AHBA and BrainSpan Atlas, we reported that these robust connectome hubs have a spatiotemporally distinctive transcriptomic pattern in contrast to non-hub regions. These results advanced our understanding of the robustness of macroscopic functional connectome hubs and their potential cellular and molecular substrates.

Extant reports have shown largely inconsistent and less reproducible hub localizations^7, 8, 13–19^, which may arise from high heterogeneity in the included subjects, data acquisition, and analysis strategies across studies. To diminish these potential confounding factors, we employed stringent participant inclusion criteria that included only healthy young adults aged 18 to 36 years and adopted harmonized data preprocessing and connectome analysis protocols across cohorts. Nevertheless, the random-effects meta-analysis revealed high heterogeneity among cohorts in almost all brain areas, which implied that heterogeneity of imaging scanners and/or imaging protocols could be an important cause for inconsistent and less reproducible results across prior studies. Thus, our study was indispensable by conducting a harmonized random-effects metaanalysis model in which both intracohort variation (i.e., sampling errors) and intercohort heterogeneity were considered^48^. In addition, our validation results showed that the spatial distribution of functional connectome hubs was relatively stable when using more than 510 subjects and 35 cohorts, demonstrating that 5,212 subjects from 61 cohorts were adequate to minimize both sampling errors and heterogeneity among cohorts. Considering only dozens of subjects in most prior studies^7, 8, 13–15, 17, 19^, the low statistical power attributed to inadequate subjects could be another cause for prior inconsistent and less reproducible hub localizations. Finally, we used harmonized image processing and connectome analysis protocols across cohort, which avoided methodological variation and reduced potential methodological defects that have not been resolved in prior studies. See an extension discussion in Supplementary Text II.

The present results demonstrated that the 35 highly consistent and reproducible connectome hubs show heterogeneous functional connectivity profiles, forming three clusters. Twenty-one hubs (Cluster I) are connected with extensive areas in the DAN, VAN, FPN, and SMN. Previous investigations indicated that they are core regions of the DAN (left AIP, right 7PC, left 7Am, bilateral PFt, left FEF, bilateral 6a, right 6v, and right FST)^29, 49^, VAN (left 43, left FOP4, right 46, right 6r, right PF, left PFop, left SCEF, right 5mv)^29, 49^, and FPN (left p9-46v and right IFSa)^29, 50^. In addition, hub regions involved in the sensorimotor pathway (right VIP, right FST, left 7Am, and left FEF)^51^ are also connected with the visual association cortex, acting as connectors between the VIS and the SMN, DAN, and VAN. Information flow along the primary visual, visual association, and higher-level sensorimotor cortices is undertaken by the four occipital hubs (Cluster II) left VMV1, right V4, and bilateral V3A that are all densely connected with the VIS and portions of the SMN, DAN, and VAN. This aligns with the role of their homologous regions in the non-human primate cerebral cortex^51^. The remaining 10 hubs (Cluster III) are all located in canonical DMN regions^52^. One of them, the left 8Av hub, is robustly connected with both DMN and lateral prefrontal FPN regions, acting as a connector between the DMN and FPN. This can be supported by the recent finding of a control-default connector located in the posterior middle frontal gyrus^32^ and may also be a case of the hypothesis of parallel interdigitated subnetworks^53^. This observation offers a significant complementary interpretation to the conventional assumption that the DMN is anticorrelated with other networks^52^. Considering that communication between the DMN and other networks is of significant relevance to neuropsychiatric disorders^54^, such as autism spectrum disorders^55^, we speculated that the left 8Av hub may be a promising target region for therapeutic interventions.

We demonstrated that these robust brain hubs have a spatiotemporally distinctive transcriptomic pattern dominated by genes with the highest enrichment for the neuropeptide signaling pathway. Because neuropeptides are a main type of synaptic transmitter that is widely distributed in the human central nervous system^56^, robust neuropeptide signaling pathways are indispensable for efficient synaptic signal transduction that sustains dense and flexible functional connections of hub regions. In addition, hub regions have higher transcription levels for main neuronal metabolic pathways in contrast to non-hubs. This is reasonable because massive synaptic activities in hub regions demand high material and metabolic costs, which is in accordance with our observation of more blood supply and higher oxidative phosphorylation and aerobic glycolysis levels in hub regions. This is also in agreement with prior observations of a tight coupling between FCS and blood supply^1, 47^.

We found that connectome hubs possess a spatiotemporally distinctive transcriptomic pattern of key neurodevelopmental processes in contrast to non-hubs. Specifically, connectome hubs have higher transcription levels for dendrite and synapse development and lower transcription levels for axon development and myelination during childhood, adolescence, and adulthood. These findings are compatible with previous observations of the prefrontal area having higher synapse density but lower myelination than primary somatosensory, auditory, and visual (V1/V2) cortices^43, 44^. Higher transcription levels for dendrite and synapse development in hub regions are necessary for the overproduction of synapses that will be selectively eliminated based on the demand of the environment and gradually stabilized before full maturation^57^, which is “the major mechanism for generating diversity of neuronal connections beyond their genetic determination”^58^. Lower transcription levels for axon development and myelination will prolong the myelination period in hub regions, which characterizes a delayed maturation phase^59^. Marked delay of anatomical maturation in human prefrontal and lateral parietal cortices has been frequently observed both in human development^58, 60^ and in primate evolution^59^, which provides more opportunities for social learning to establish diverse neuronal circuits that contribute to our complex^58^ and species-specific^59^ cognitive capabilities. We also observed higher transcription levels for neuron migration in hub regions from mid-fetal period to early infancy. This is in agreement with the report of extensive migration of young neurons persisting for several months after birth in the human frontal cortex^61^. Meanwhile, the migration and final laminar positioning of postmitotic neurons are regulated by common transcription factors^62^, which suggests that a higher transcription level for neuron migration in hub regions facilitates the construction of more intricate interlaminar connectivity. These microscale divergences of key neurodevelopmental processes may result in a more intricate macroscale structural connectivity proflie in hub regions.

Human neurodevelopment is an intricate and protracted process, during which the transcriptome of the human brain requires precise spatiotemporal regulation^38^. Thus, in addition to contributing to our complex cognitive capabilities, the distinctive transcriptomic pattern of neurodevelopment in hub regions may also increase connectome hubs’ susceptibility to neuropsychiatric disorders^58, 59^, which means small disturbance in the magnitude or the timing of this transcriptomic pattern may have long-term consequences on brain anatomical topography or functional activation. This is in line with our observation of psychiatric disorders being the most significant disease associated with the top 150 key genes (Table S5). This implies that uncovering the intricate transcriptomic pattern, diverse neuronal circuits, anatomical topography, and functional activation of connectome hubs provide crucial and promising routes for understanding the pathophysiological mechanisms underlying neurodevelopmental disorders, such as autism spectrum disorders^38, 55^ and schizophrenia^5, 38, 58, 59^.

Of note, we conducted transcriptome-connectome association analysis using machine learning approaches in which non-linear mathematical operations were implemented rather than linear operations, such as linear correlation^24^, linear regression^25^, or partial least squares^26^. It has been argued that observations of transcriptome-connectome spatial association have a high falsepositive rate through linear regression^63^ and linear correlation^64^ and may be largely shifted toward the first principal component axis of the dataset through partial least squares^65^. These investigations imply that prior transcriptome-connectome association results by linear mathematical operations may include high false-positive observations that are independent of connectome measurements, such as genes enriched for ion channels^24–26^. By contrast, high reproducibility across different machine learning models and across different GO enrichment analysis tools and convergent results from the AHBA and BrainSpan Atlas made it very unlikely that our findings were false-positive observations.

Some results of the present study should be interpreted cautiously because of methodological issues. First, we identified the robust connectome hubs using preprocessed rsfMRI data with global signal regression because of its great promise in minimizing physiological artifacts on functional connectomes^66^. Validation analysis demonstrated that hub distribution identified without global signal regression was more likely derived from physiological artifacts rather than by ongoing neuronal activity (Supplementary Text and Fig S9). Second, the AHBA dataset only includes partial human genes, of which approximately half were excluded in data preprocessing^34^, which may have induced incomplete observations in our data-driven analysis. Finally, our transcriptomic signature results addressed only the association between connectome hubs and transcriptomic patterns and did not explore causation between them. Exploring more detailed mechanisms underlying this association is attractive and may be practicable for nonhuman primate brains in future studies.

## Methods

### Dataset

We included a large-sample rsfMRI dataset of 5,212 healthy young adults (aged 18–36 years, 2,377 males) across 61 cohorts from Asia, Europe, North America, and Australia. Data of each cohort were collected with participants’ written informed consent and with approval by the respective local institutional review boards. All data passed strict quality controls and were routinely preprocessed with a uniform pipeline. For details, see Supplementary Text I.

### Identifying robust functional connectome hubs using a harmonized meta-analysis

For each individual, we constructed a voxelwise functional connectome matrix by computing the Pearson’s correlation coefficient between preprocessed rsfMRI time series of all pairs of voxels within a predefined gray matter mask (47,619 voxels). The gray matter mask was divided into seven large-scale cortical networks^29^ and a subcortical network^30^. The cerebellum was not included due to largely incomplete coverage during rsfMRI scanning in most cohorts. Negative functional connections were excluded from our analysis due to neurobiologically ambiguous interpretations^67^. To further reduce signal noise and simultaneously avoid potential sharing signals between nearby voxels, both weak connections (Pearson’s *r* < 0.1) and connections terminating within 20 mm were set to zero^68^. We validated the threshold of weak connections using 0.05 and 0.2.

For each voxel, we computed the FCS as the sum of connection weights between the given voxel and all the other voxels. We further normalized this resultant FCS map with respect to its mean and standard deviation across voxels^7^. For each cohort, we performed a general linear model on these normalized FCS maps to reduce age and gender effects. For each voxel, we constructed the general linear model as:

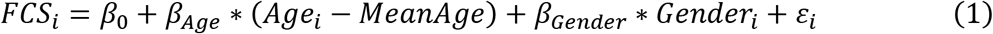

*FCS_i_, Age_i_, Gender_i_*, and *ε_i_* indicate the FCS, age, gender, and residual of the *i*th individual, respectively. *MeanAge* indicates the mean age of that cohort. The general linear model exported a mean FCS map and its corresponding variance map for each cohort.

The mean and variance FCS maps of the 61 cohorts were submitted to a random-effects metaanalysis model^48^ to address across-cohort heterogeneity of functional connectomes. The detailed computational procedures are described in the book^48^. A short summary of these procedures was provided in Supplementary Text I. This resulted in a consistent FCS pattern and its corresponding SE map. We compared the FCS of each voxel with the average of the whole brain (i.e., zero) using a *Z* value^48^:

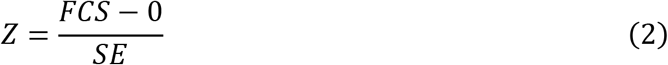

In line with previous neuroimaging meta-analysis study^69^, we performed 10,000 one-sided nonparametric permutation tests^28^ to assign a *p* value to the observed *Z* value. For each iteration, after randomizing the spatial correspondence among cohorts’ mean FCS maps (the spatial correspondence between a cohort’s mean FCS map and its variance map was not changed), we repeated the computation procedure of the random-effects meta-analysis for each voxel and extracted the maximum *Z* value of all voxels to construct a null distribution. A *p* value was assigned to each voxel by comparing the observed *Z* value to the null distribution. For a significance level below 0.05, this *p* value closely tracks the Bonferroni threshold^28^. Finally, we defined functional connectome hubs as brain regions with a *p* value less than 0.001 and cluster size greater than 200 mm^3^. The thresholds of *p* value and cluster size were similar with the activation likelihood estimation algorithm^69^. We extracted MNI coordinates for each local peak *Z* value terminating beyond 15 mm within each brain cluster using the *wb_command -volume-extrema* command (https://humanconnectome.org/software/workbench-command/-volume-extrema) in Connectome Workbench v1.4.2. Effect size was estimated using Cohen’s *d* metric ^48^:

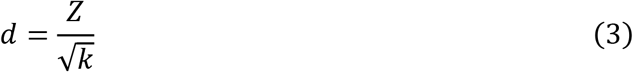

*k* is the number of cohorts in the meta-analysis.

### Mapping seed-to-whole-brain connectivity profiles of functional connectome hubs

We modeled each hub seed region as a sphere with a 6-mm radius centered on the hub peak and computed Pearson’s correlation coefficients between the seed region’s preprocessed rsfMRI time series and the time series of all gray matter voxels. The time series of the seed region was computed by averaging the time series of all gray matter voxels in the seed sphere. These correlation coefficients were further transformed to Fisher’s *z* for normality. In line with above, we constructed a general linear model on these Fisher’s *z* value maps within each cohort to reduce age and gender effects and performed a random-effects meta-analysis on these Fisher’s *z* value maps across cohorts to address the across-cohort heterogeneity, resulting in a robust Fisher’s *z* pattern and its corresponding SE map. Then, We compared the Fisher’s *z* value of each voxel with zero using a *Z* value^48^ and estimated effect size using Cohen’s *d* metric^48^ as described in equations (2) and (3). We performed 10,000 one-sided nonparametric permutation tests^28^ to identify the most consistent functional connection *Z* value map with a *p* value less than 0.001 and cluster size greater than 200 mm^3^. Finally, we divided the connectivity map of each hub into eight brain networks mentioned above and represented the functional connectivity profile of a hub as the voxel percentage of each of the eight networks connected with it to address the effect of network size. To illustrate the left 8Av hub’s connectivity profile, we also mapped its homologous region the right 8Av region’s connectivity profile (Fig S5).

### Identifying transcriptomic signatures underlying functional connectome hubs

We trained classifiers based on XGBoost and SVM to distinguish connectome hubs from non-hubs using transcriptomic features from the preprocessed AHBA dataset^34^. The top 150 genes (Table S1) mostly contributed to the classification results were submitted to GO enrichment analyses using GOrilla^35^ (http://cbl-gorilla.cs.technion.ac.il) and DAVID^36, 37^ v6.8 (https://david.ncifcrf.gov). For details, see Supplementary Text I. Based on GO enrichment analysis results, we tested transcription level differences of gene sets involved in key neurodevelopmental processes^38^ (Table S6) and main neuronal metabolic pathways^39^ (oxidative phosphorylation^40^ and aerobic glycolysis^41^, Table S7) between connectome hubs and non-hubs through one-sided Wilcoxon rank-sum test. In line with prior studies^38, 41^, we used the first principal component of each gene set’s transcription level to plot and to perform the statistical analysis (Fig 5A). For illustration purposes, we normalized the first principal component of each gene set’s transcription level respect to its minimum and maximum values across all brain samples to range from 0 to 1.

To explore developmental details, we inspected transcriptomic trajectory differences between connectome hubs and non-hubs in the above gene sets using the BrainSpan Atlas^42^. In line with prior studies^38, 41^, we used the first principal component of each gene set’s transcription level to plot transcriptomic trajectories and visually inspected transcriptomic trajectory differences between connectome hubs and non-hubs (Fig 5B). Transcriptomic trajectories were plotted using locally weighted regression by smoothing the first principal component of each gene set’s transcription level against log2[post-conceptional days] as in prior study^38^. Of note, considering apparent transcriptomic differences compared to the neocortex^38^, we excluded the striatum, mediodorsal nucleus of the thalamus, and cerebellar cortex in the transcriptomic trajectory analysis but not the amygdala and hippocampus whose transcriptomic trajectories are more similar to those of the neocortex than to those of other subcortical structures^38^. Analysis using only neocortical regions revealed almost unchanged results (Fig S8).

To validate above results derived from transcriptome datasets, we tested fiber number differences between connectome hubs and non-hubs through one-sided Wilcoxon rank-sum test (Fig 6). Fiber number data across different length bins was derived from a fiber length profiling dataset^45^. For each fiber length bin, fiber number of each vertex was normalized with respect to its mean and standard deviation across vertices. We further examined differences between connectome hubs and non-hubs in metabolic measurements of blood supply (the cerebral blood flow), oxidative phosphorylation (the cerebral metabolic rate for oxygen), and aerobic glycolysis (the glycolytic index) through one-sided Wilcoxon rank-sum test (Fig 7). These measurements were derived from a positron emission tomography study^46^ and assigned to 82 Brodmann areas and seven subcortical structures. Brodmann areas with more than 50% vertices or subcortical structures with more than 50% voxels identified as hubs were regarded as hub regions.

### Statistical analysis

We performed statistical analyses using MATLAB R2013a. Statistical significance of brain clusters in Fig 2B and 3A and Fig S2B, S3B, S5, and S9A were determined by comparing the observed *Z* values in equation (2) with their corresponding null distributions constructed by above mentioned 10,000 one-sided nonparametric permutation tests^28^. To determine the statistical significance of one-sided Wilcoxon rank-sum tests in Fig 5A, 6, and 7 and Fig S9D, we constructed 1,000 surrogate hub identification maps with the spatial autocorrelations being corrected using a generative model^70^ and repeated calculating rank-sum statistics using these surrogate hub identification maps to construct a null distribution. Then, *p* values derived by comparing the observed rank-sum statistics with their corresponding null distributions were Bonferroni-corrected. Surrogate hub identification maps in Fig 5A, 6, and 7 were constructed based on the hub identification map in Fig 2B. Surrogate hub identification maps in Fig S9D were constructed based on the hub identification map in Fig S9A.

## Data availability

The MRI data of the first 60 cohorts listed in Table S8 are available at the International Neuroimaging Data-sharing Initiative (http://fcon_1000.projects.nitrc.org), Brain Genomics Superstruct Project (https://doi.org/10.7910/DVN/25833), Human Connectome Project (https://www.humanconnectome.org), MPI-Leipzig Mind-Brain-Body Project (https://openneuro.org/datasets/ds000221), and Age-ility Project (https://www.nitrc.org/projects/age-ility). The MRI data of the PKU cohort are under active use by the reporting laboratory and will be available upon reasonable request. The preprocessed AHBA dataset is available at https://doi.org/10.6084/m9.figshare.6852911. The normalized BrainSpan Atlas dataset is available at http://brainspan.org/static/download.html. The fiber length profiling dataset^45^ is available at https://balsa.wustl.edu/study/1K3l.

## Code availability

The code to reproduce the results and visualizations of this manuscript is available at https://github.com/zhileixu/FunctionalConnectomeHubs.

## Author Contributions

Conceptualization: Z.X., Y.H.; Methodology: Z.X., Y.H., M.X., X.W., X.L., T.Z.; Investigation: Z.X.; Visualization: Z.X.; Supervision: Y.H.; Writing—original draft: Z.X., Y.H.; Writing— review & editing: Y.H., Z.X., M.X., X.L., T.Z., X.W.

## Acknowledgments

We thank Drs. Huali Wang and Xiaodan Chen for data acquisition of the PKU cohort and Drs. Qiushi Wang and Nan Zhang for valuable advice on MRI data quality controls. This work was supported by the National Natural Science Foundation of China [82021004, 31830034, and 81620108016 to Y.H., 82071998 and 81671767 to M.X., 81971690 to X.L., 81801783 to T.Z.], Changjiang Scholar Professorship Award [T2015027 to Y.H.], National Key Research and Development Project [2018YFA0701402 to Y.H.], Beijing Nova Program [Z191100001119023 to M.X.], and Fundamental Research Funds for the Central Universities [2020NTST29 to M.X.].

