## Supplementary Text for "Mapping consistent, reproducible, and transcriptionally relevant functional connectome hubs of the human brain"

Supplementary Information: 20 text pages, 9 figures, 10 tables.

### Supplementary Text I

**Image preprocessing and quality control.**

We collected a large-sample rsfMRI dataset ($N$ = 7,202) from public data-sharing platforms and in-house cohorts, which consists of 73 cohorts from Asia, Europe, North America, and Australia. Data of each cohort were collected with participants' written informed consent and with approval by the respective local institutional review boards.

We first reviewed T1-weighted structural MRI data for all participants with the assistance of a neuroradiologist and a clinical neurologist to confirm no identifiable lesion or structural abnormality (e.g., regional atrophy and posterior cranial fossa arachnoid cyst). All rsfMRI data for the remaining participants were preprocessed routinely using SPM12 v6470 (https://www.fil.ion.ucl.ac.uk/spm/software/spm12) and GRETNA v2.0.0[1] with a uniform pipeline. For each individual, we discarded the first 10s' volumes for magnetic field stabilization and the participant's adaptation to the scanner. Next, slice-timing was corrected within each volume. To correct for head motion, all volumes were realigned to the mean image. Participants with significant head motion (translation above 3 mm or rotation above 3° in any direction) were excluded from the subsequent analyses. Then, all volumes were normalized to the 3-mm isotropic space of the Montreal Neurological Institute (MNI) using the EPI template provided by SPM12. The normalized volumes were spatially smoothed using a 6-mm full-width at half-maximum Gaussian kernel. After that, the time series of each voxel underwent the procedure of linear trend removal, nuisance signal regression (24 head motion parameters, white matter, cerebrospinal fluid, and global brain signals), and temporal band-pass filtering (0.01-0.1 Hz). Finally, scrubbing was performed to minimize head motion effects[2]. Specifically, volumes with framewise displacement exceeding 0.5 mm and their adjacent volumes (1 back and 2 forward) were replaced with linearly interpolated data. We excluded participants with more than 25% interpolated volumes. Notably, slice-timing was not corrected in the Human Connectome Project cohort due to multiband acquisition[3]. For participants with more than one rsfMRI scan, we used only one of them. To reduce the potential effects of development and aging on our results, we restricted our analysis to healthy young adults (aged 18–36 years). To ensure sufficient statistical power, twelve cohorts were discarded due to having fewer than 10 participants that passed quality controls. After these stringent quality controls, we included preprocessed rsfMRI data of 5,212 healthy young adults (2,377 males) from 61 cohorts in the final analysis. The sample size and age ranges of each cohort were summarized in Fig 1. Table S8 provides detailed information on the individual cohorts.

**Random-effects meta-analysis.**

To deal with both intracohort variation (i.e., sampling errors) and intercohort heterogeneity, we adopted a random-effects meta-analysis model[4] to obtain the consistent pattern across cohorts. The detailed computational procedures of the random-effects meta-analysis model are described in the book[4]. Here, we give a short summary of these procedures.

For each voxel, $M_i$, $SD_i$, and $N_i$ indicate the mean and standard division value and the participant number of the $i$th cohort, respectively. The original weight assigned to the $i$th cohort is the inverse of its variance:

$$W_i = \frac{N_i}{{SD_i}^2} \tag{1}$$

The heterogeneity between cohort means was calculated as:

$$Q = \sum W_i {M_i}^2 - \frac{(\sum W_i M_i)^2}{\sum W_i} \tag{2}$$

The expected value of $Q$ is the degrees of freedom:

$$df = k - 1 \tag{3}$$

where $k$ is the number of cohorts in the meta-analysis. Therefore, the estimated variance of the cohort mean distribution was calculated as:

$$T^2 = \frac{Q - df}{\sum W_i - \frac{\sum {W_i}^2}{\sum W_i}} \tag{4}$$

The percentage of total variability that reflects heterogeneity among cohorts was calculated as:

$$I^2 = \frac{Q - df}{Q} \times 100\% \tag{5}$$

The weight assigned to the $i$th cohort was updated as:

$$W_i^* = \frac{1}{\frac{{SD_i}^2}{N_i} + T^2} \tag{6}$$

The result of the random-effects meta-analysis was calculated as:

$$M^* = \frac{\sum {W_i}^* M_i}{\sum {W_i}^*} \tag{7}$$

The variance of $M^*$ was estimated as:

$$V_{M^*} = \frac{1}{\sum {W_i}^*} \tag{8}$$

The standard error of $M^*$ was calculated as:

$$SE_{M^*} = \sqrt{V_{M^*}} \tag{9}$$

This random-effects meta-analysis model exported a mean value $M^*$, its corresponding standard error value $SE_{M^*}$, and the heterogeneity score $I^2$.

**Identifying transcriptomic signatures underlying functional connectome hubs.**

*AHBA dataset.* The original AHBA consists of microarray expression data of more than 20,000 genes in 3702 spatially distinct brain samples taken from six neurotypical adult donors[5]. Only two out of six donors were sampled from both hemispheres and the other four were sampled from only the left hemisphere. Because “no statistically significant hemispheric differences could be identified”[5], a prior study[6] provided a publicly available preprocessed AHBA dataset that includes 10,027 genes’ transcriptomic data for 1,285 left cortical samples. Preprocessing steps taken by the study[6] mainly include probe-to-gene re-annotation, intensity based data filtering, probe selection, accounting for individual variability, and gene filtering. We downloaded this preprocessed AHBA dataset form https://doi.org/10.6084/m9.figshare.6852911. Of the 1,285 samples, 382 were identified as hub samples and 776 as non-hub samples according to their MNI coordinates and the hub identification map in Fig 2B. The remaining 127 samples were not included in our analysis because they were out of our gray matter mask. The brain samples used in our analysis are listed in Table S9.

*BrainSpan dataset.* The normalized BrainSpan dataset provided by previous study[7] was downloaded from http://brainspan.org/static/download.html. It was generated using 524 brain samples from 42 donors aged from eight post-conceptional weeks to 40 postnatal years, including 52,376 genes’ transcriptomic data for 11 neocortical areas and five additional regions of the human brain. The brain regions used in our analysis are listed in Table S10.

*XGBoost classifier.* We built a supervised classifier based on XGBoost, a scalable tree boosting system with state-of-the-art resource efficiency and superior performance in many machine learning challenges[8], to distinguish hub samples from non-hub samples using 10,027 genes’ transcriptomic data from the preprocessed AHBA dataset. We used equal amounts of positive samples (hub samples) and negative samples (non-hub samples) in the classifier training procedure to ensure that the optimal classifier was unbiased toward any type of sample. To balance the time complexity and the prediction accuracy, we trained the classifier with 300 randomly selected hub samples and 300 randomly selected non-hub samples and tested it with the remaining 82 hub samples and 476 non-hub samples. Based on previous experience[9], we performed a 30-fold cross-validation procedure to identify the optimal number of model training iterations with the following parameters: *nrounds* = 1,500, *early_stopping_rounds* = 50, *eta* = 0.05, *objective* = “binary:logistic”. The sensitivity, specificity, and accuracy rate of the XGBoost classifier were stably estimated by repeating the randomly selecting training samples, cross-validation, and classifier training and testing procedures 1,000 times. We implemented XGBoost using the *XGBoost* package[8] v1.2.0.1 in R 4.0.2.

*SVM classifier.* To exclude the XGBoost model’s potential bias relating to mostly contributed key genes, we reproduced classification results using another machine learning model based on SVM. Instead of using all 10,027 genes’ transcriptomic features, we only used genes with the greatest contributions to the XGBoost classifier to train the SVM classifier. If the key genes with the greatest contributions to the classification results are independent of the XGBoost model, the SVM classifier would achieve a comparable or higher accuracy rate than the XGBoost classifier, even though the SVM classifier was trained using only partial genes’ transcriptomic features. In line with the XGBoost model, we used equal amounts of hub samples and non-hub samples in the classifier training procedure. For the easiest classification task, the SVM classifier was trained to distinguish all 382 hub samples from 382 non-hub samples with the highest rate to be correctly classified by the XGBoost classifier. For the most difficult classification task, the SVM classifier was trained to distinguish all 382 hub samples from 382 non-hub samples with the lowest rate to be correctly classified by the XGBoost classifier. To balance the time complexity and the prediction accuracy, we performed a leave-two-out cross-validation procedure to obtain the optimal SVM classifier. We implemented SVM using the *svm* function from the *scikit-learn* package[10] v0.23.2 in Python 3.8.3 with default parameters.

*GO enrichment analysis using GOrilla.* We conducted two GO enrichment analyses using GOrilla[11] (http://cbl-gorilla.cs.technion.ac.il). The first analysis used the 150 key genes with the greatest contributions to the XGBoost classifier as the target list and all 10,027 genes as the background list. The second analysis used the ranked 10,027 genes according to their contributions to the XGBoost classifier. Of note, we performed GO enrichment analyses for all three ontology categories: biological process, molecular function, and cellular component. However, only analysis for biological process yielded significant GO terms.

*GO enrichment analysis using DAVID.* We repeated GO enrichment analysis for biological process using DAVID[12, 13] v6.8 (https://david.ncifcrf.gov) with the 150 key genes with the greatest contributions to the XGBoost classifier as the target list and all 10,027 genes as the background list. In addition, we also performed GO enrichment analysis for disease association using the 150 key genes as the target list and all 10,027 genes as the background list.

### Supplementary Text II

**Methodological variation or defects may cause controversial hub reports.**

Methodological variation or defects in prior studies may cause controversial hub reports in specific regions, such as the primary areas, subcortical structures, and cerebellum. The controversy may arise from multifaceted sources.

Both unimodal and primary areas have been argued as candidate hub regions[14-19]. Our results demonstrated that functional connectome hubs were located in many unimodal cortices but not in the primary cortex. This finding is supported by the functional organization of the human cerebral cortex, where the primary cortex confines functional connectivity mostly in the primary cortex and portions of the unimodal cortex, whereas the unimodal cortex bridges the primary and heteromodal cortices[20, 21]. Prior study demonstrated that the primary cortex possesses most short-range functional connections[18], which could be systematically but spuriously elevated by subject head motion[2] or by unavoidable signal blurring across sulci or gyri[22]. These spurious short-range functional connections can be effectively suppressed by motion scrubbing, global signal regression, and stringent participant inclusion criteria as well as excluding short-range correlations[2, 22, 23], which all have been thoroughly addressed in the present study but were largely neglected in prior reports. Accordingly, it is reasonable to speculate that functional connectome hubs in the primary cortex reported in prior studies should be driven by spurious short-range functional connections.

Most subcortical structures have also been argued as candidate hubs, including the thalamus[15, 16, 24], basal ganglia[15, 25], amygdala[16, 25], and hippocampus[25]. Nevertheless, no subcortical structure was identified as a candidate hub in the present study. The inconsistency may be attributed to more complex sampling error and intercohort heterogeneity in subcortical structures. First, a prior report[26] demonstrated reliable estimation of subcortical-cortical functional connections requiring more data (~100 min per subject) than conventional quantities of rsfMRI data (5–20 min per subject) adopted by prior reports[15, 16, 24, 25]. In addition, individual features contribute to ~60% of the variance in subcortical-cortical functional connections[26], which is higher than ~35% in cortical-cortical[27] and ~45% in cerebellar-cortical[28] functional connections. It precludes reliable estimation of subcortical functional connections with only dozens of subjects. These two factors complicate both sampling error and intercohort heterogeneity in subcortical structures, which is in line with our observation of higher heterogeneity among cohorts in most subcortical structures than in the cortex (Fig S1).

A portion of the cerebellum has also been argued as a candidate hub region[15, 16, 25]. Although the cerebellum was not included in the present study due to largely incomplete coverage during rsfMRI scanning in most cohorts, these reports conflicted with the metabolic pattern of the human brain. The cerebellum has been demonstrated to have the lowest level of aerobic glycolysis and a lower metabolic rate for glucose than most cortical regions[29], which makes it unfeasible for the cerebellum to maintain and run dense functional connections during rest. Combined with previous findings, we speculate that hub reports in the cerebellum should be caused by inexplicable negative functional connections[25], spurious short-range functional connections[15, 16], or unstable estimation of functional connections due to inadequate data[28].

Functional connectome hubs have been reported in the superior temporal gyrus[14, 17-19, 25, 30], but rarely in the rolandic operculum. Our results demonstrated the superior temporal gyrus with subequal FCS levels compared with the rolandic operculum (Fig 2A). However, we identified significant hub peaks in the rolandic operculum and surrounding regions, such as the left area 43 and right 6r, rather than in the superior temporal gyrus. This finding may result from rsfMRI signal blurring across sulci because of the close proximity of the superior temporal gyrus and rolandic operculum[20]. Signal blurring may result in potential hub peaks in the superior temporal gyrus merging with those in the rolandic operculum and surrounding regions, causing false-negative observations of hub peaks in the superior temporal gyrus. Nevertheless, signal blurring is unavoidable in rsfMRI data processing, such as realignment, resampling, registration, and subject averaging[22]. Consequently, limited data resolution complicates the interpretation of functional connectome hubs in the superior temporal gyrus and rolandic operculum. This issue should be resolved in future studies using rsfMRI data with higher spatial resolution and greater signal specificity.

A prior study argued that identifying hub regions based on FCS highlights only members of large brain networks rather than brain regions playing crucial roles in global brain communication[22]. However, no significant correlation between a voxel's FCS and the size of the brain network to which it belongs could be identified in the present study (Fig S4). The conclusion in the study[22] may be driven by unreasonable connection threshold (Pearson's *r*: 0.20-0.37) because we observed all hub peaks possessing significant connections with Pearson's *r* less than 0.2 (Fig S7).

### Supplementary Text III

**Global signal effect on spatial distribution of functional connectome hubs.**

To examine the effect of global signal regression on hub distribution, we repeated identifying functional connectome hubs using preprocessed rsfMRI data without global signal regression. Fig S9A shows that hub distribution was largely shifted by rsfMRI data preprocessing without global signal regression, of which some canonical hubs were absent, such as the inferior parietal gyrus, anterior insula, medial prefrontal cortex, and post cingulate cortex; more middle and anterior cingulate, visual, and auditory cortices were identified as hubs. This shifted hub distribution corresponds well with the report of brain regions with a global signal correlation significantly higher than the average[31]. It is compatible with our analysis of the global signal localization (GSL) score, which localizes the spatial distribution of the global signal (Fig S9B). Specifically, for each individual, we computed the Fisher's $z$ transformed Pearson's correlation coefficient between the global signal and preprocessed rsfMRI time series of each voxel. Next, we constructed a general linear model on these Fisher's z value maps within each cohort to reduce age and gender effects and performed a random-effects meta-analysis on these Fisher's z value maps across cohorts to address the across-cohort heterogeneity, resulting in a consistent GSL map (Fig S9B). We observed that brain regions with GSL scores greater than 0.5 significantly overlapped with the shifted hub distribution (*Dice* = 0.833, $p < 0.001$, Fig S9C).

Considering prior observations of a tight coupling between FCS and blood supply[32, 33], we examined differences between connectome hubs and non-hubs in metabolic measurements of blood supply (the cerebral blood flow) using the hub distribution in Fig S9A. But one-sided Wilcoxon rank-sum test shown no significant difference between connectome hubs and non-hubs in the cerebral blood flow ($p = 0.077$, Fig S9D).

Together, the hub distribution identified using preprocessed rsfMRI data without global signal regression was more likely derived from physiological artifacts rather than by the intrinsic or ongoing neuronal activity.

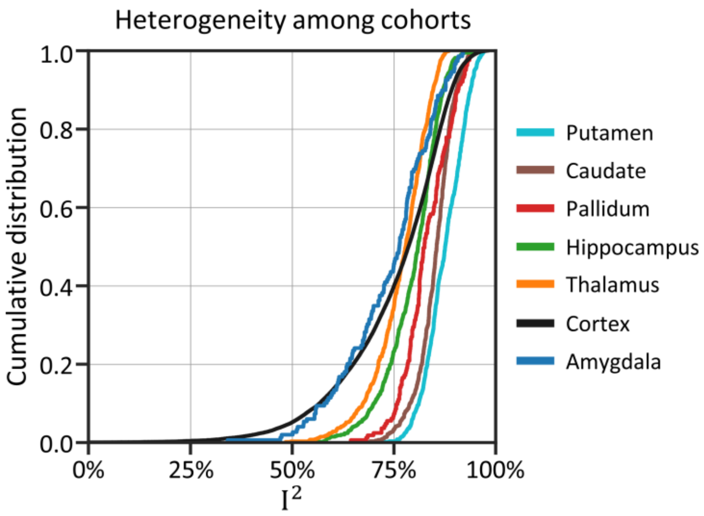


**Fig S1. Cumulative distribution function plot of $I^2$.**

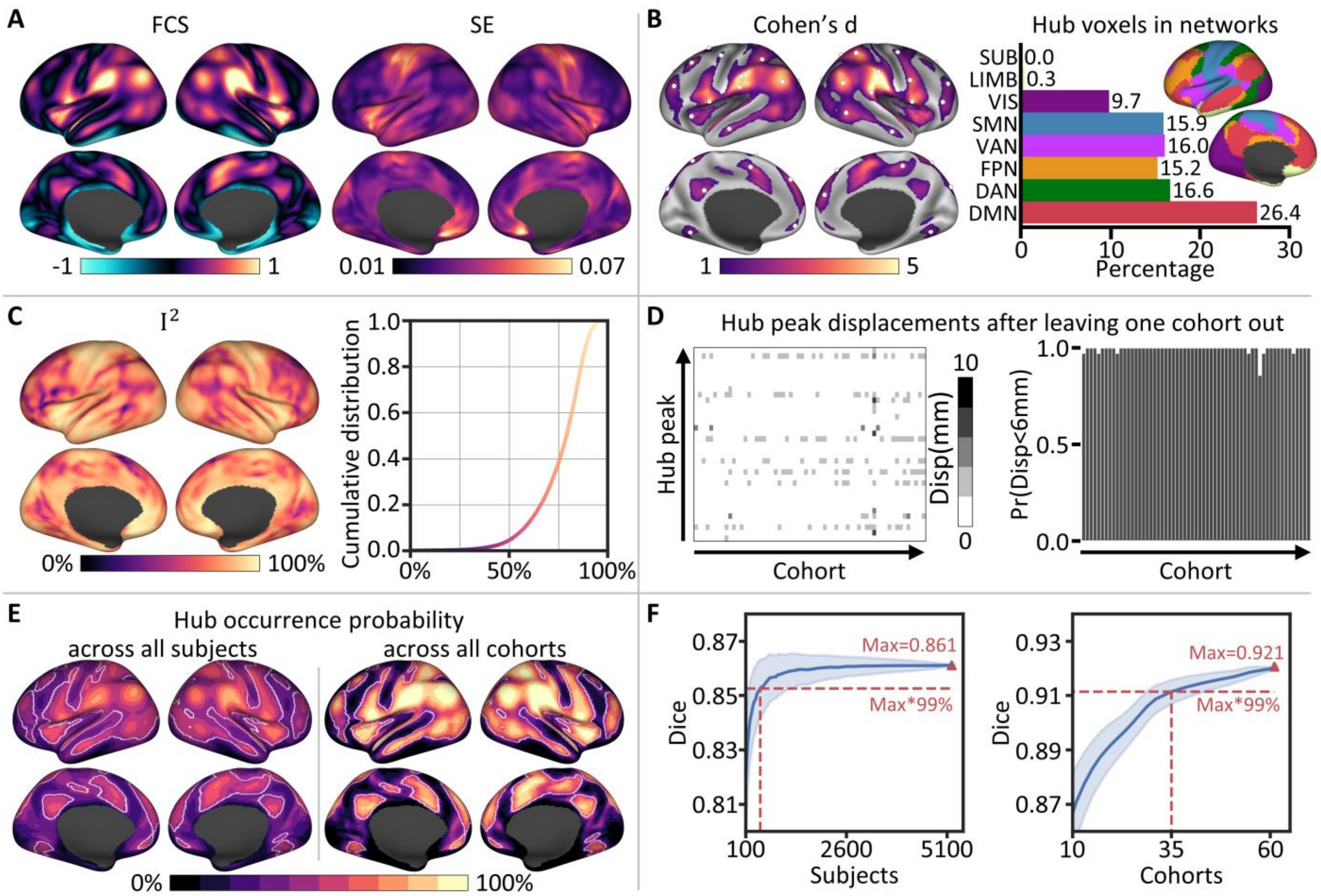


**Fig S2. Highly consistent and reproducible functional connectome hubs using a connection threshold of 0.05.** **A** Robust FCS pattern and its corresponding variance (standard error, SE) map estimated using a harmonized voxelwise random-effects meta-analysis across 61 cohorts. **B** Left: The most consistent functional connectome hubs ($p < 0.001$, cluster size > 200 $mm^3$); white spheres represent hub peaks. Right: Hub voxel distribution in eight large-scale brain networks; insets, the seven large-scale cortical networks[20] were rendered on the left hemisphere. **C** Left: Heterogeneity measurement $I^2$ estimated through the random-effects meta-analysis. Right: Cumulative distribution function plot of $I^2$. **D** Left: Heatmap of displacements of the 35 hub peaks after leaving one cohort out. Right: Bar plot of the probability across the 35 hub peaks whose displacement was less than 6 mm after leaving one cohort out. **E** Hub occurrence probability map across all subjects (left) and all cohorts (right). White lines delineate boundaries of the identified hubs in **B**. **F** Dice's coefficient of the identified hubs in **B** compared with the top *N* (voxel number of the identified hubs in **B**) voxels with the highest hub occurrence probability values across randomly selected subjects (left) and randomly selected cohorts (right). Blue shading represents the standard deviation across 2,000 random selections.

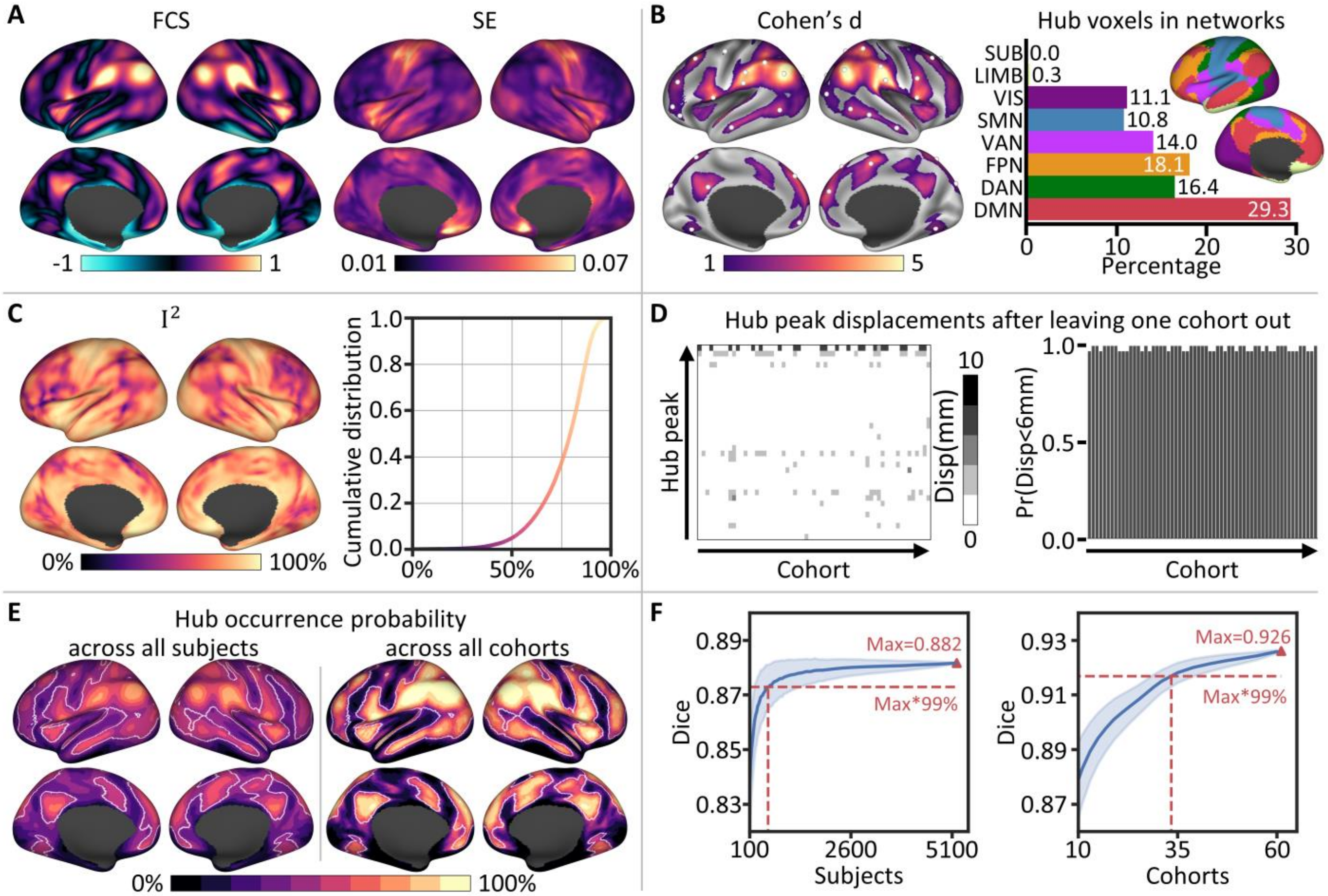


**Fig S3. Highly consistent and reproducible functional connectome hubs using a connection threshold of 0.2.** **A** Robust FCS pattern and its corresponding variance (standard error, SE) map estimated using a harmonized voxelwise random-effects meta-analysis across 61 cohorts. **B** Left: The most consistent functional connectome hubs ($p < 0.001$, cluster size > 200 $mm^3$); white spheres represent hub peaks. Right: Hub voxel distribution in eight large-scale brain networks; insets, the seven large-scale cortical networks[20] were rendered on the left hemisphere. **C** Left: Heterogeneity measurement $I^2$ estimated through the random-effects meta-analysis. Right: Cumulative distribution function plot of $I^2$. **D** Left: Heatmap of displacements of the 35 hub peaks after leaving one cohort out. Right: Bar plot of the probability across the 35 hub peaks whose displacement was less than 6 mm after leaving one cohort out. **E** Hub occurrence probability map across all subjects (left) and all cohorts (right). White lines delineate boundaries of the identified hubs in **B**. **F** Dice's coefficient of the identified hubs in **B** compared with the top *N* (voxel number of the identified hubs in **B**) voxels with the highest hub occurrence probability values across randomly selected subjects (left) and randomly selected cohorts (right). Blue shading represents the standard deviation across 2,000 random selections.

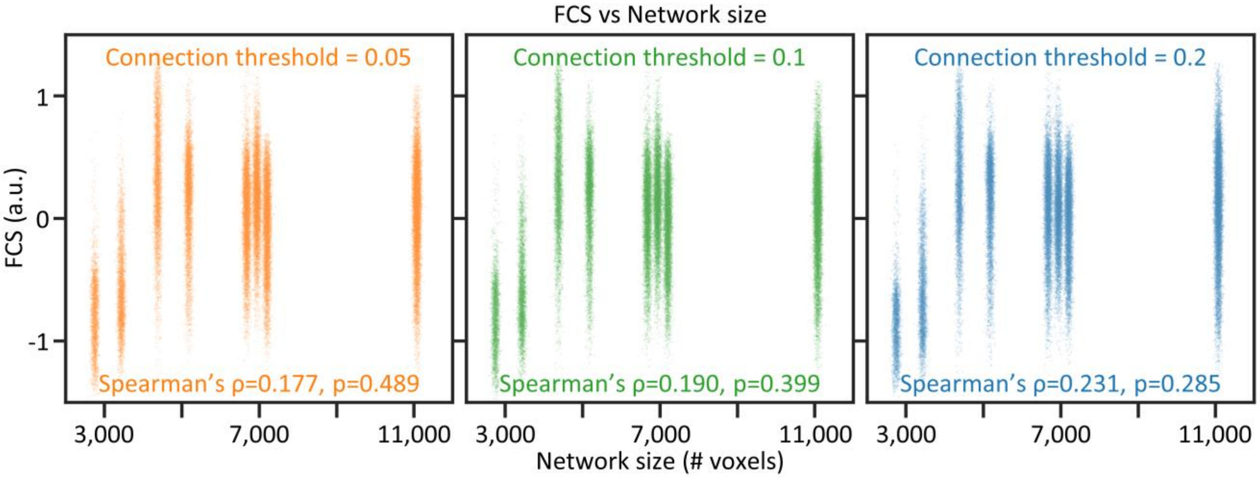


**Fig S4. Relationship between FCS and network size.** Scatter plot showing no significant correlation between the FCS of voxels and the size of the brain network to which they belong. The *p* value was estimated through a nonparametric permutation test with 10,000 iterations and Bonferroni-corrected. For each iteration, the voxel number of each network was reshuffled. Each dot represents a voxel. For illustration purposes, dots were jittered along the x axis (uniform jitter of ±50 voxels). a.u., arbitrary unit.

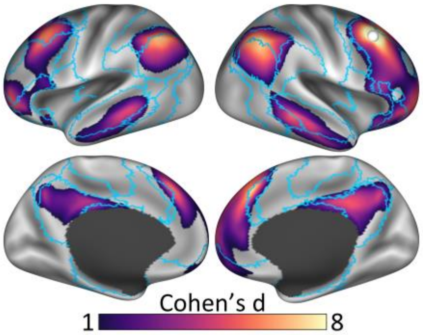


**Fig S5. Functional connectivity profile of the right 8Av region.** White spheres represent the right 8Av seed (MNI coordinates: 45, 18, 45). Blue lines delineate boundaries of the seven cortical networks shown in Fig 2B.

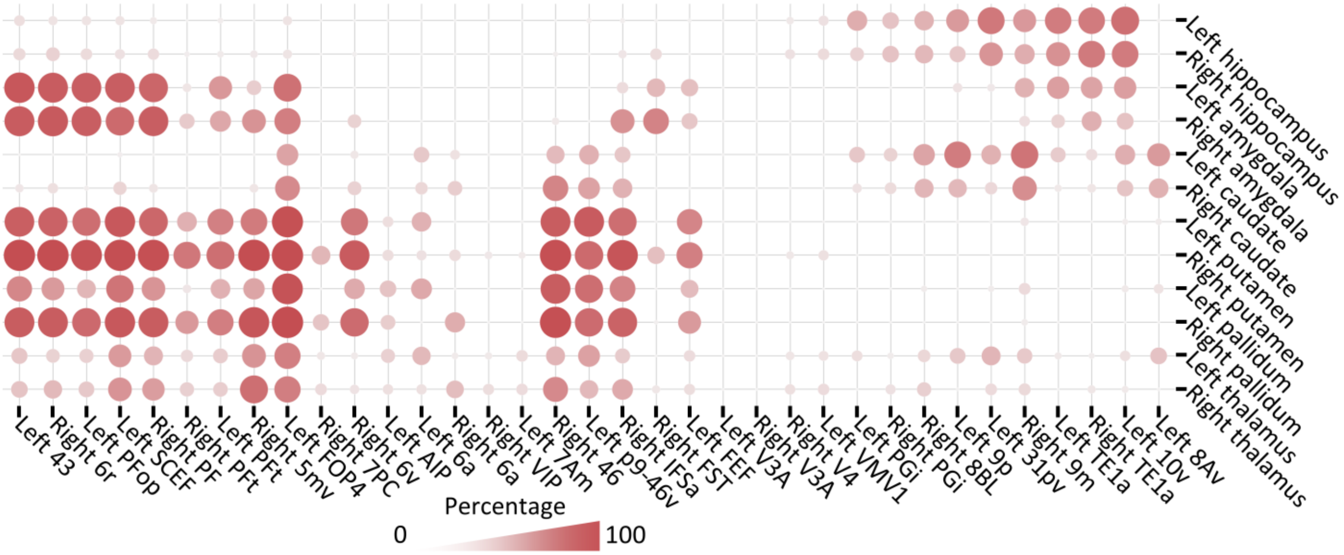


**Fig S6. Brain hubs' connectivity profiles with subcortical nucleus.** Percentage matrix showing brain hubs' heterogeneous connectivity profiles with subcortical nucleus. Each item of the percentage matrix represents the voxel percentage of one subcortical nucleus connected with one hub.

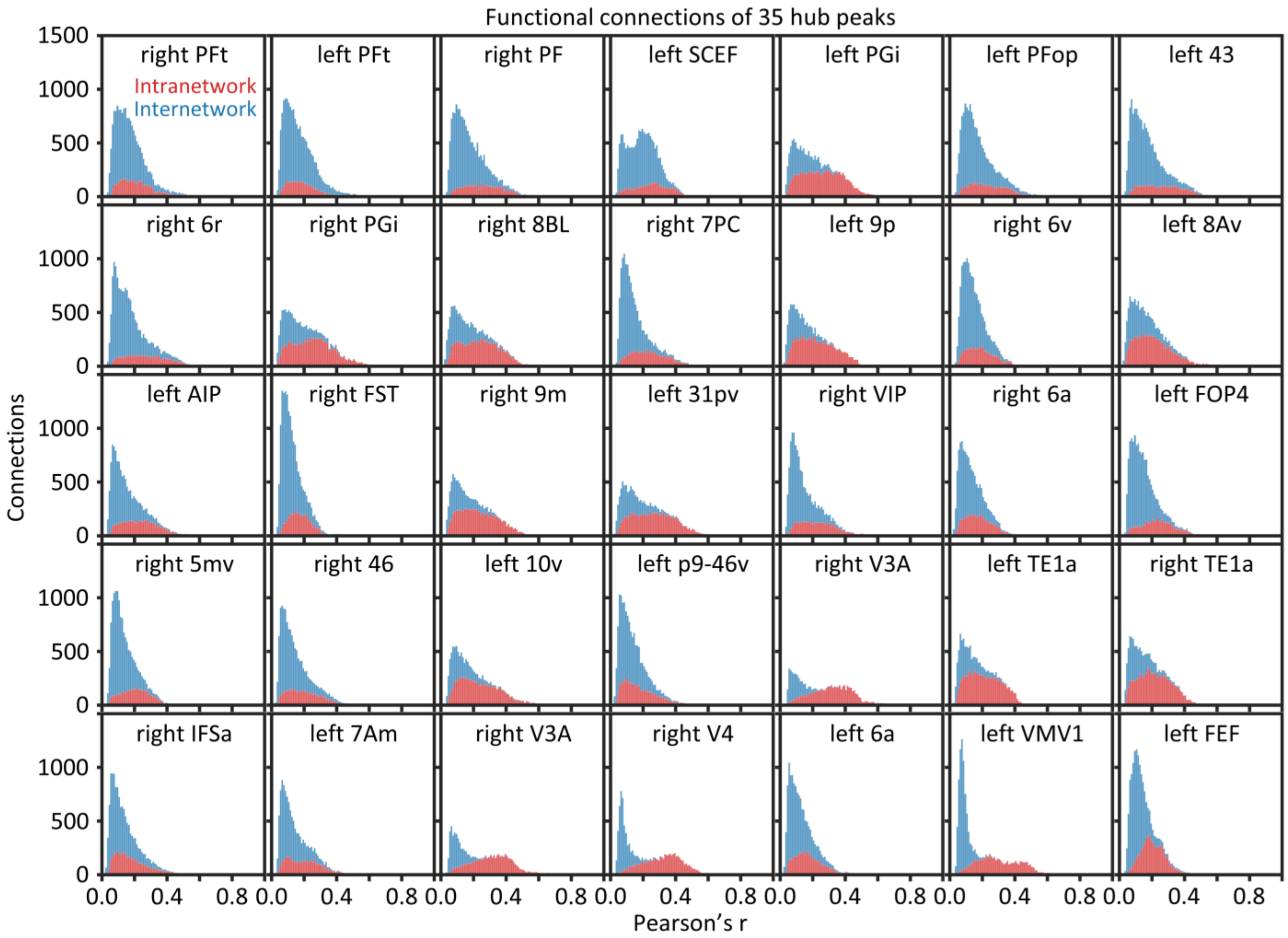


**Fig S7. Histogram plot of the connection strength of each hub's robust functional connectivity profile shown in Fig 3A.** Intranetwork and iternetwork connections are displayed as stacked.

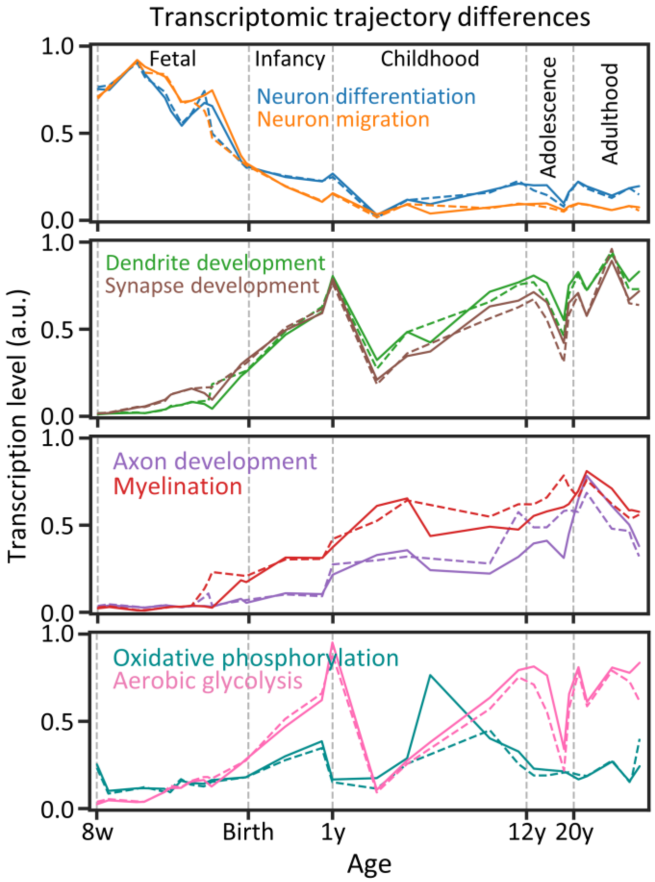


**Fig S8. Transcriptomic trajectory analysis using only neocortical regions.** w, post-conceptional week; y, postnatal year; a.u., arbitrary unit.

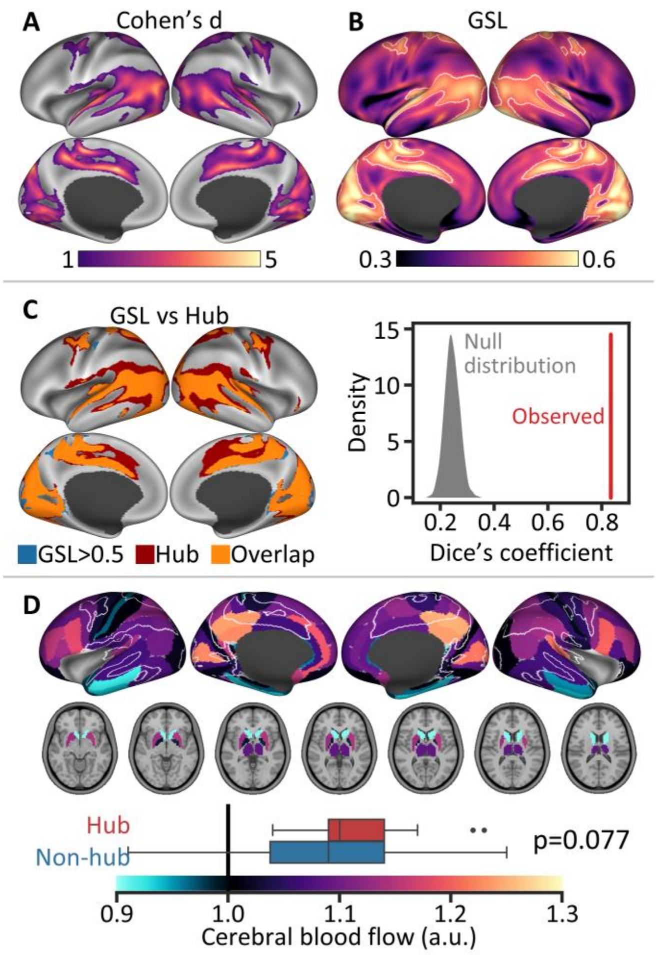


**Fig S9. Global signal effect on spatial distribution of functional connectome hubs. A** Functional connectome hubs identified using preprocessed rsfMRI data without global signal regression. Brain regions with FCS significantly higher than zero were identified as hubs ($p < 0.001$, cluster size > 200 $mm^3$). **B** Global signal localization (GSL) score distribution. Brain regions with GSL scores greater than 0.5 were delineated with white lines. **C** Overlap of brain regions with GSL scores greater than 0.5 on the identified hubs in **A**. The null distribution of the Dice's coefficient was constructed by generating 1,000 surrogate map of the hub distribution map in **A** with the spatial autocorrelations being corrected using a generative model[34]. **D** The cerebral blood flow of 82 Brodmann areas and seven subcortical structures were provided by a prior study[29]. White lines delineate boundaries of the identified hubs in **A**. Boxplot edges, gray lines, and whiskers and dots depict the 25th and 75th percentiles, median, and extreme nonoutlier and outlier values, respectively. Brodmann areas with more than 50% vertices or subcortical structures with more than 50% voxels identified as hubs were regarded as hub regions (n=21), vice versa as non-hub regions (n=68). Significance of one-sided Wilcoxon rank-sum test was determined by 1,000 permutation tests and were labeled with Bonferroni-corrected *p* values. a.u., arbitrary unit.

Tables S1 to S10 are provided in a separate xlsx file.

Table S1. Genes' contributions to the XGBoost classifier.

Table S2. GOrilla GO enrichment analysis results for the top 150 key genes.

Table S3. GOrilla GO enrichment analysis results for the ranked 10,027 genes.

Table S4. DAVID GO enrichment analysis results for the top 150 key genes.

Table S5. DAVID disease association analysis results for the top 150 key genes.

Table S6. Genes associated with key neurodevelopment processes.

Table S7. Genes associated with main neuronal metabolic pathways.

Table S8. Included cohorts in the final analysis.

Table S9. Included AHBA samples in the final analysis.

Table S10. Included BrainSpan samples in the final analysis.
